# Alcohol inhibits sociability via serotonin inputs to the nucleus accumbens

**DOI:** 10.1101/2023.05.29.542761

**Authors:** Ruixiang Wang, Kanza M. Khan, Nagalakshmi Balasubramanian, Thomas James, Selvakumar Govindhasamy Pushpavathi, David Kim, Samantha Pierson, Qi Wu, Mark J. Niciu, Marco M. Hefti, Catherine A. Marcinkiewcz

## Abstract

Social interaction is a core component of motivational behavior that is perturbed across multiple neuropsychiatric disorders, including alcohol use disorder (AUD). Positive social bonds are neuroprotective and enhance recovery from stress, so reduced social interaction in AUD may delay recovery and lead to alcohol relapse. We report that chronic intermittent ethanol (CIE) induces social avoidance in a sex-dependent manner and is associated with hyperactivity of serotonin (5-HT) neurons in the dorsal raphe nucleus (DRN). While 5-HT^DRN^ neurons are generally thought to enhance social behavior, recent evidence suggests that specific 5-HT pathways can be aversive. Using chemogenetic iDISCO, the nucleus accumbens (NAcc) was identified as one of 5 regions that were activated by 5-HT^DRN^ stimulation. We then employed an array of molecular genetic tools in transgenic mice to show that 5-HT^DRN^ inputs to NAcc dynorphin neurons drive social avoidance in male mice after CIE by activating 5-HT_2C_ receptors. NAcc dynorphin neurons also inhibit dopamine release during social interaction, reducing the motivational drive to engage with social partners. This study reveals that excessive serotonergic drive after chronic alcohol can promote social aversion by inhibiting accumbal dopamine release. Drugs that boost brain serotonin levels may be contraindicated for individuals with AUD.

## Main

Alcohol use disorder (AUD) afflicts an estimated 1.4% of the global population, but despite its prevalence, few effective treatments exist. Individuals with AUD may initially use alcohol to self-medicate for social anxiety and social phobia, but these symptoms tend to worsen with continued alcohol use^1^ and lead to social isolation, which is a risk factor in alcohol dependency and relapse^2–5^. Selective serotonin reuptake inhibitors (SSRIs) are still widely used in the treatment of co-morbid mood and anxiety symptoms in individuals with AUD, but not all individuals respond to this treatment, and some may experience a worsening of their symptoms, which can elevate the risk of relapse. Serotonin (5-HT) is classically thought to have mood- boosting effects, but accumulating evidence suggests that enhancing systemic serotonin can be aversive in certain contexts depending on which serotonin pathways are activated, the relative distribution of 5-HT receptors, and activity in other neural circuits that interact with serotonin^6–9^. Evidence from human and rodent literature also supports the idea that serotonin may play an important role in the aversive aspects of chronic alcohol use. Tryptophan hydroxylase-2, the biosynthetic enzyme for 5-HT biosynthesis, is upregulated in individuals with AUD^6, 10^, while studies in rodents suggest that dorsal raphe nucleus (DRN) serotonergic neurons are more active during alcohol withdrawal^11, 12^. Here we used DREADD-assisted 3D imaging to identify serotonergic projections that may be implicated in alcohol-induced social deficits, and we further employed pathway-specific viral-genetic manipulations to dissect the specific circuit mechanisms driving social avoidance in the context of AUD.

### Sex-dependent effects of ethanol withdrawal on social behavior

We first assessed social deficits in adult male and female C57BL/6J mice with a history of chronic intermittent ethanol (CIE). These mice were given voluntary access to two-bottle choice ethanol for 8 weeks, which resulted in an average ethanol intake and preference of 13.09 ± 2.36 g/kg/24-h and 46.8 ± 8.9%, respectively, for males and 26.12 ± 4 g/kg/24-h and 57± 6% for females at the 8-week mark (Extended Data Figure 1A-B). Despite lower ethanol consumption and preference overall, male mice exhibited social deficits in the social interaction test (Bonferroni t_36_=7.33, p<0.0001) after CIE whereas females did not. Females also spent less time in social interaction relative to males (Bonferroni t_36_=3.41, p<0.01) (Extended Data Figure 1C-E). No anxiety-like behaviors or locomotor deficits were observed in the elevated plus maze (Extended Data Figure 1F-I). In the sucrose preference test, there was no 3-way CIE x sex x time interaction, but there was a two-way CIE x sex interaction (F_1,36_=4.44, p<0.05) as well as a main effect of sex (F_1,36_=19.82, p<0.0.0001) and time (F_3.81, 136.4_=2.72, p<0.05), but not CIE (Extended Data Figure 1J). There was also no change in time spent in the center of an open field in either sex, but males exhibited hypoactivity in the open field which may be indicative of anxiety (Bonferroni t_36_=3.20, p<0.05) (Extended Data Figure 1K-O). Since the ethanol-induced social deficits were limited to males, we used male mice throughout the remainder of the study.

**Figure 1:**
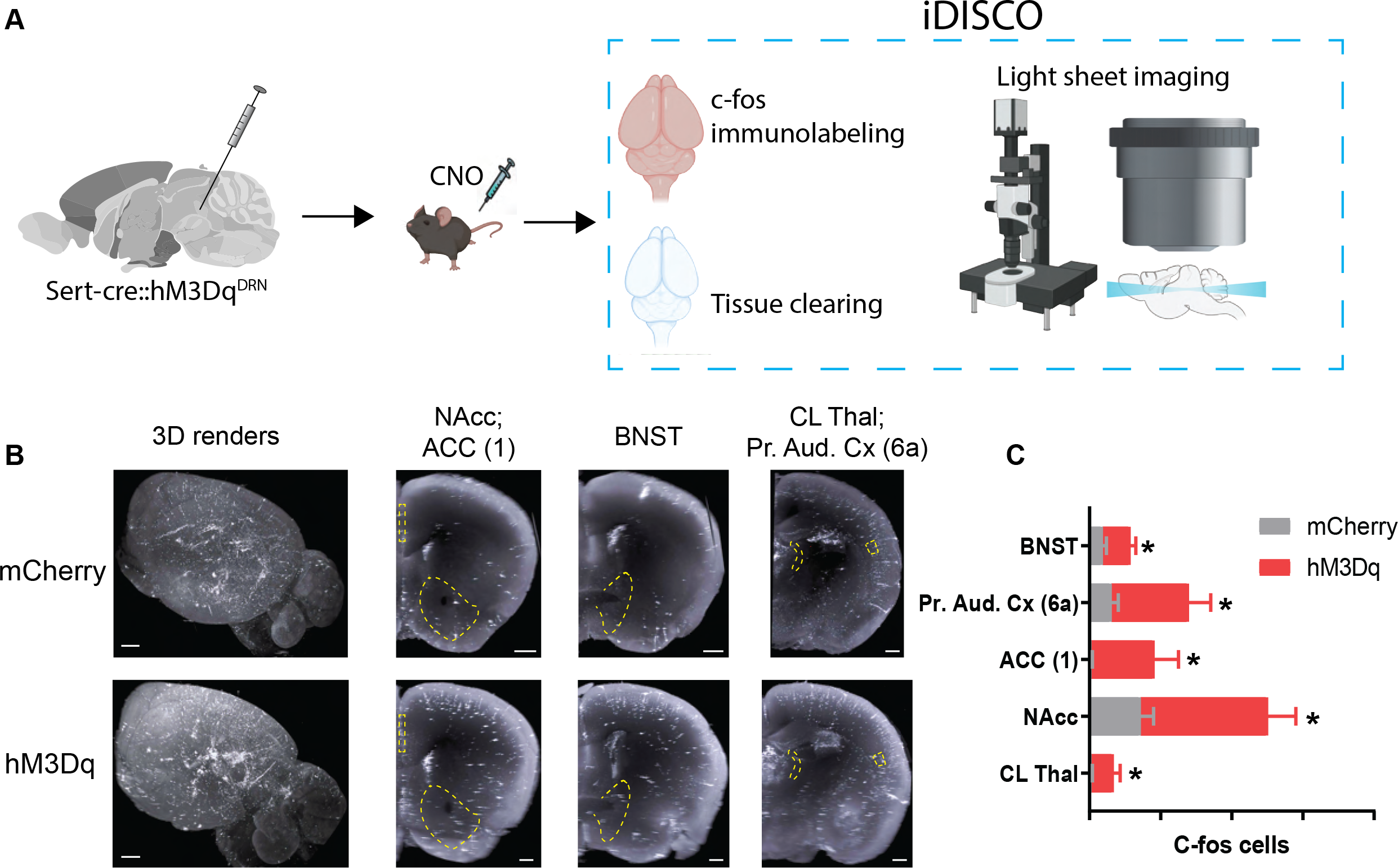
DREADD-assisted mapping of activated outputs of 5-HT^DRN^ neurons. (A) Experimental configuration of viral infusions into Sert-cre mice and CNO induction of c-fos expression, immunolabeling and tissue clearing, and light sheet imaging. (B) 3D rendering of c-fos expression in a representative control (mCherry) and hM3Dq-injected mouse (scale bars = 700 µm), and coronal views through the NAcc, ACC (layer 1), BNST, primary auditory cortex (layer 6a), and central lateral thalamus (scale bars = 500 µm). (C) Histogram of c-fos-positive cell counts in select brain regions. * p<0.05. *Parts of this figure were made using Biorender*.

### Mapping 5-HT^DRN^-stimulated brain-wide activation patterns with chemogenetic iDISCO

Since 5-HT^DRN^ neurons have been linked to social behavior^13^, we first asked whether CIE could alter the excitable properties of these neurons. *Ex vivo* electrophysiological recordings revealed that 5-HT neurons in CIE mice had lower mean rheobase (i.e. action potential threshold) (t_15_=2.73, p<0.05) and increased current-induced spiking which is indicative of enhanced excitability (Interaction F_20,280_=3.57, p<0.0001; Area under the curve: t_14_=4.02, p<0.01) (Extended Data Figure 2). We then paired chemogenetic stimulation of 5-HT^DRN^ neurons with whole brain c-fos immunolabeling and volumetric imaging (iDISCO)^14^ to map neuronal activity in downstream targets. *Sert-cre* mice were stereotaxically injected with a Cre-inducible Gq-coupled DREADD (hM3Dq) in the DRN, and their brains were harvested for iDISCO 3 hours after CNO injection. Volumetric images were aligned to the Allen Brain Atlas and c-fos-immunoreactive (IR) cells were counted using the ClearMap spot detection algorithm. C-fos-IR neurons were elevated in the central lateral nucleus of the thalamus (CL thalamus), nucleus accumbens (NAcc), ventral anterior cingulate cortex (ACC, layer 1), primary auditory cortex (layer 6a), and bed nucleus of the stria terminalis (BNST) (Figure 1). The NAcc has been previously implicated in social behavior in the context of autism spectrum disorder and morphine withdrawal^15, 16^, so we focused on the 5-HT projections to the NAcc throughout the remainder of the study.

**Figure 2:**
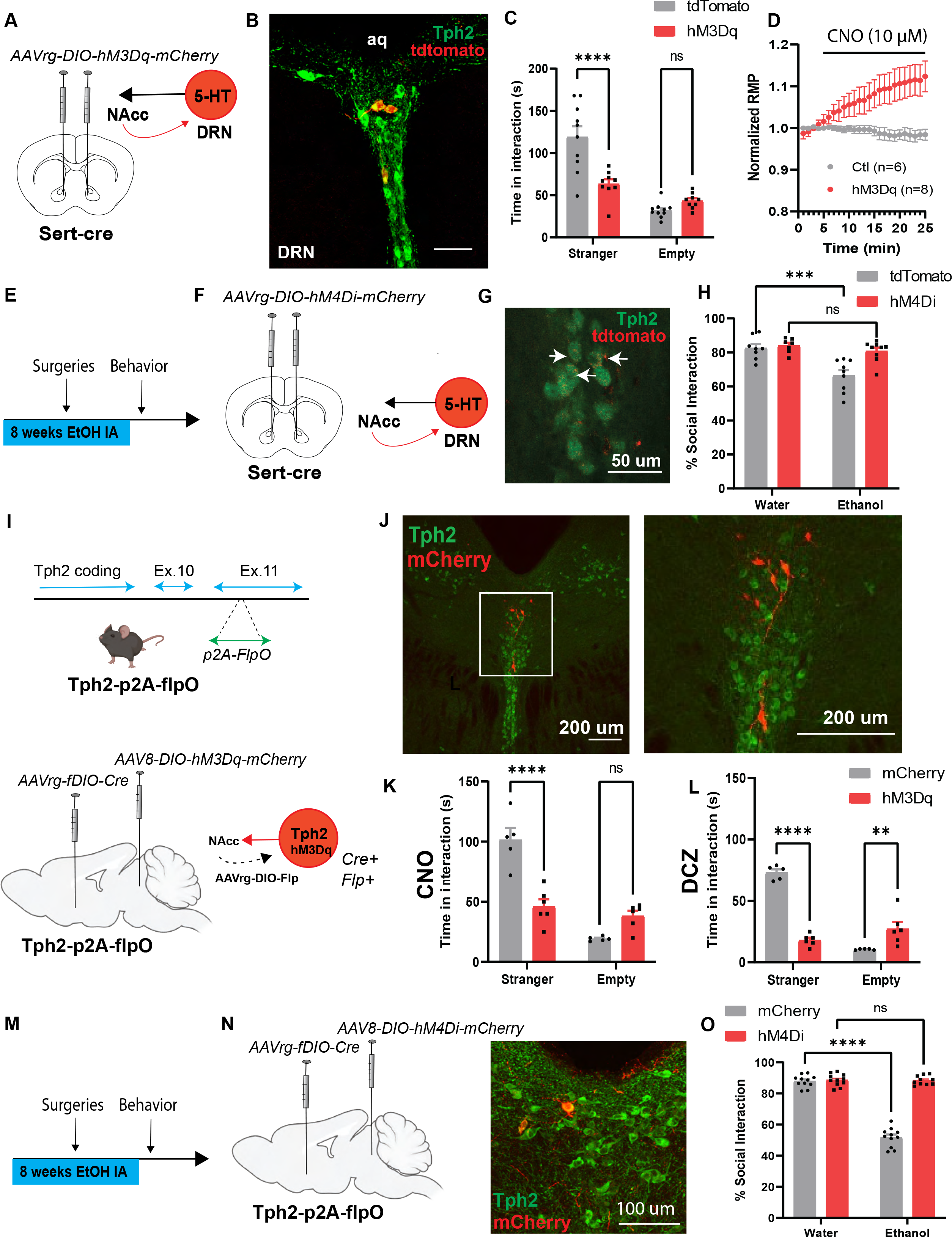
5-HT^DRN^**^è^**^NAcc^ neurons drive CIE-induced social deficits. (A) Experimental schematic of viral targeting of Gq-coupled DREADDs to NAcc-projecting 5-HT neurons. (B) Confocal image of hM3Dq-expressing 5-HT neurons in the DRN (scale bar = 50 µm; aq: aqueduct). (C) Chemogenetic activation of NAcc-projecting 5-HT neurons induces social deficits in ethanol-naïve mice. (D) Functional validation of DREADDs in electrophysiological recordings from 5-HT^DRN^ neurons showing a positive voltage deflection during bath application of 10 µM CNO. RMP: resting membrane potential. (E) Experimental schematic of chronic intermittent access to ethanol, and (F) viral targeting of Gi-coupled DREADDs to NAcc projecting 5-HT^DRN^ neurons. (G) Confocal images of hM4Di-expressing 5-HT^DRN^ neurons (scale bar = 50 µm). (H) Chemogenetic inhibition of NAcc-projecting 5-HT neurons mitigates social deficits after CIE. (I) Targeting vector used to generate Tph2-p2A-flpO mice (upper panel) and schematic depicting AAV-mediated intersectional targeting of Gq-coupled DREADDs to 5-HT^DRNèNAcc^ neurons (lower panel). (J) Confocal images showing hM3Dq-mCherry expression in 5-HT^DRNèNAcc^ neurons (scale bars = 200 µm). (K-L) Chemogenetic activation of 5-HT^DRNèNAcc^ neurons with CNO (3 mg/kg, i.p.) or DCZ (100 µg/kg, i.p.) induces social deficits in ethanol-naïve mice. (M) Experimental timeline of CIE, surgeries, and behavioral tests. (N) Viral targeting of Gi- coupled DREADDs to 5-HT^DRNèNAcc^ neurons and confocal image of DREADD expression in 5-HT^DRNèNAcc^ neurons (scale bar = 100 µm). (O) Chemogenetic inhibition of the 5-HT^DRNèNAcc^ pathway rescues social deficits in CIE mice. ** p<0.01, *** p<0.001, **** p<0.0001, ns: non-significant. *Parts of this figure were made using Biorender*.

### Neuronal activity in NAcc-projecting 5-HT neurons drives social deficits after CIE

We next targeted Gq-coupled DREADDs to NAcc-projecting 5-HT neurons by injecting ethanol-naïve *Sert-cre* mice with the retrograde virus AAVrg-hsyn-DIO-hM3Dq-mCherry or a control virus (AAVrg-hsyn-DIO-mCherry) in the NAcc. *Sert-cre*::hM3Dq^5-HT➔NAcc^ mice spent significantly less time in social interaction relative to controls after CNO (F_1,17_=21.73, p<0.001; Bonferroni t_34_=5.26, p<0.0001) (Figure 2A-C). There were no group differences in anxiety-like behavior in the open field, although there was a trend toward a reduction in locomotor activity, which may be indicative of increased anxiety (Extended Data Figure 3). We then validated DREADD function in organotypic slices of the DRN by showing that bath application of CNO depolarized hM3Dq-expressing 5-HT neurons (Figure 2D). Interestingly, chemogenetic inhibition also rescued social deficits in CIE-exposed *Sert-cre*::hM4Di^5-HT➔NAcc^ mice (Interaction: F_1,29_=6.75, p<0.05, Bonferroni post-tests: Water-mCherry vs. CIE-mCherry, t_29_=4.71, p<0.001; Water- hM4Di vs. CIE-hM4Di, t_29_=0.93, ns), suggesting that increased neuronal activity in NAcc-projecting 5-HT neurons is necessary for the expression of social deficits after CIE (Figure 2E- H).

**Figure 3:**
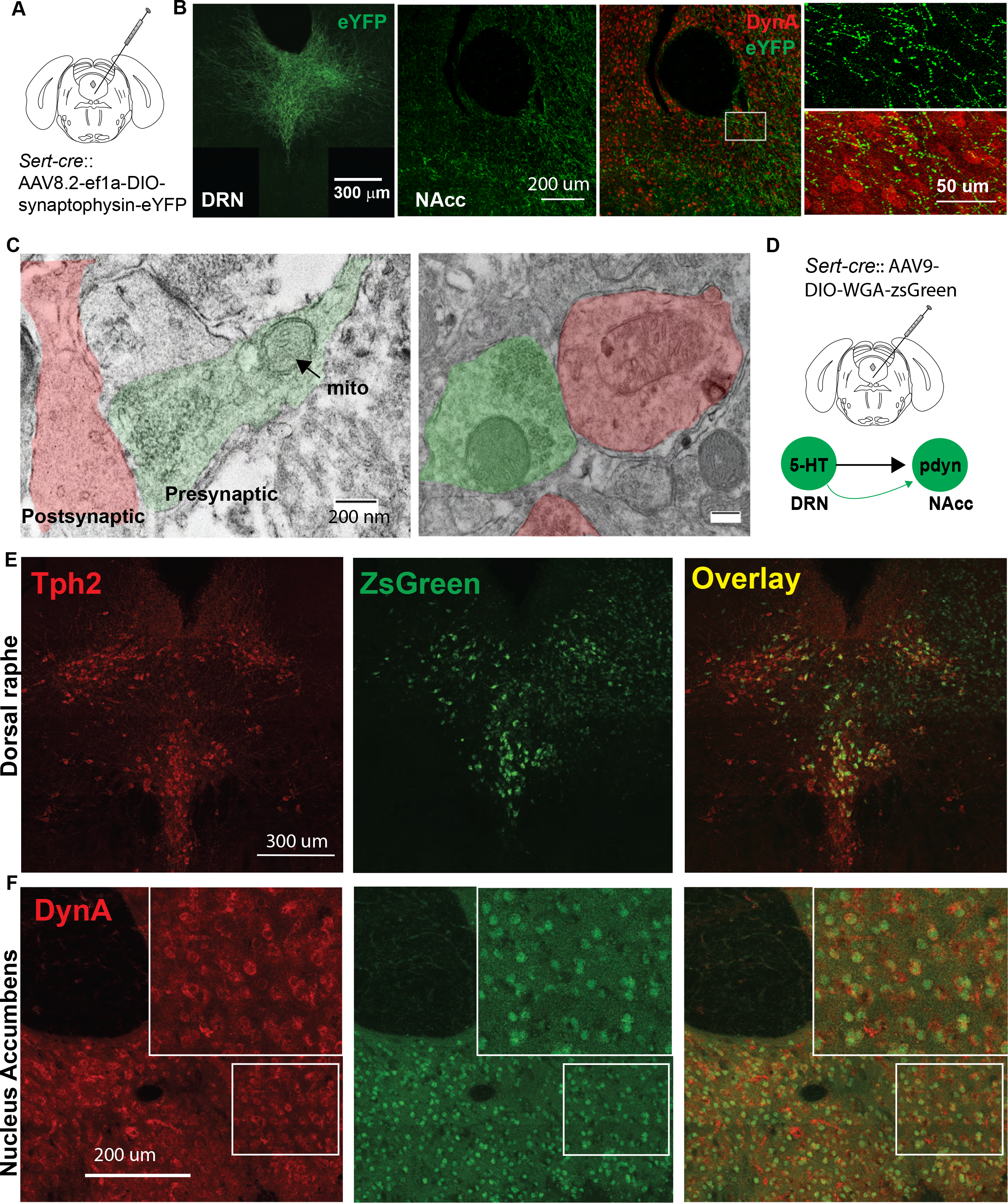
Serotonin neurons in the DRN have direct synaptic input to NAcc dynorphin neurons. (A) Experimental schematic of stereotaxic infusions of Cre-dependent synaptophysin-eYFP into the DRN. (B) Confocal images of synaptophysin-eYFP in the DRN and NAcc showing proximity of 5-HT axonal terminals to dynorphin neurons in the ventral NAcc (scale bars left to right: 300 µm, 200 µm, 50 µm). (C) Electron micrographs of synaptic junctions between eYFP-positive presynaptic terminals and dynorphin A-positive postsynaptic terminals in the ventral NAcc (scale bar = 200 nm). Mito: mitochondria. (D) Viral infusion of AAV expressing a Cre-dependent anterograde transsynaptic tracer WGA-ZsGreen in the DRN. (E) Confocal images of Tph2 (left), ZsGreen (center), and Overlay between Tph2 and ZsGreen (Right) in the DRN (scale bars = 300 µm). (F) Confocal images of dynorphin A (left), ZsGreen (center), and overlay between dynorphin A and Zsgreen (right) in the NAcc indicating a significant degree of colocalization between the two signals (scale bars = 200 µm).

This retrograde viral targeting strategy results in DREADD expression in all serotonergic nuclei that project to the NAcc, including the median raphe nucleus (MRN). To target 5-HT projections from the DRN specifically, we used a combined Cre/flp strategy. Tph2-p2A-flpO mice were generated at the University of Iowa Genome Editing Core Facility and validated in our lab by fluorescence *in situ* hybridization (FISH) and immunofluorescence colocalization of Flp-dependent mCherry with Tph2-IR neurons in the DRN (Extended Data Figure 4). We then stereotaxically injected these Tph2-p2A-flpO mice with AAVrg-fDIO-Cre in the NAcc and AAV8- DIO-hM3Dq-mCherry or a control virus (AAV8-DIO-mCherry) in the DRN to target 5-HT^DRN➔NAcc^ neurons. Chemogenetic activation of these neurons with CNO (F_1,9_=42.38, p<0.001, Bonferroni t_18_=6.57, p<0.0001) or the selective DREADD agonist deschloroclozapine (DCZ)^17^ (F_1,9_=137.4, p<0.0001, Bonferroni t_18_=11.52, p<0.0001), induced social deficits in ethanol-naïve mice (Figure 2I-L). We then employed a similar strategy to target Gi-coupled DREADDs to 5-HT^DRN➔NAcc^ neurons in Tph2-p2A-flpO mice and demonstrate that chemogenetic inhibition of this 5-HT subpopulation rescued social deficits after CIE (Interaction: F_1,39_=180, p<0.0001, Bonferroni post-tests: Water-mCherry vs. CIE-mCherry, t_39_=19.34, p<0.0001; Water-hM4Di vs. CIE-hM4Di, t_39_=0.12, ns) (Figure 2M-O).

**Figure 4:**
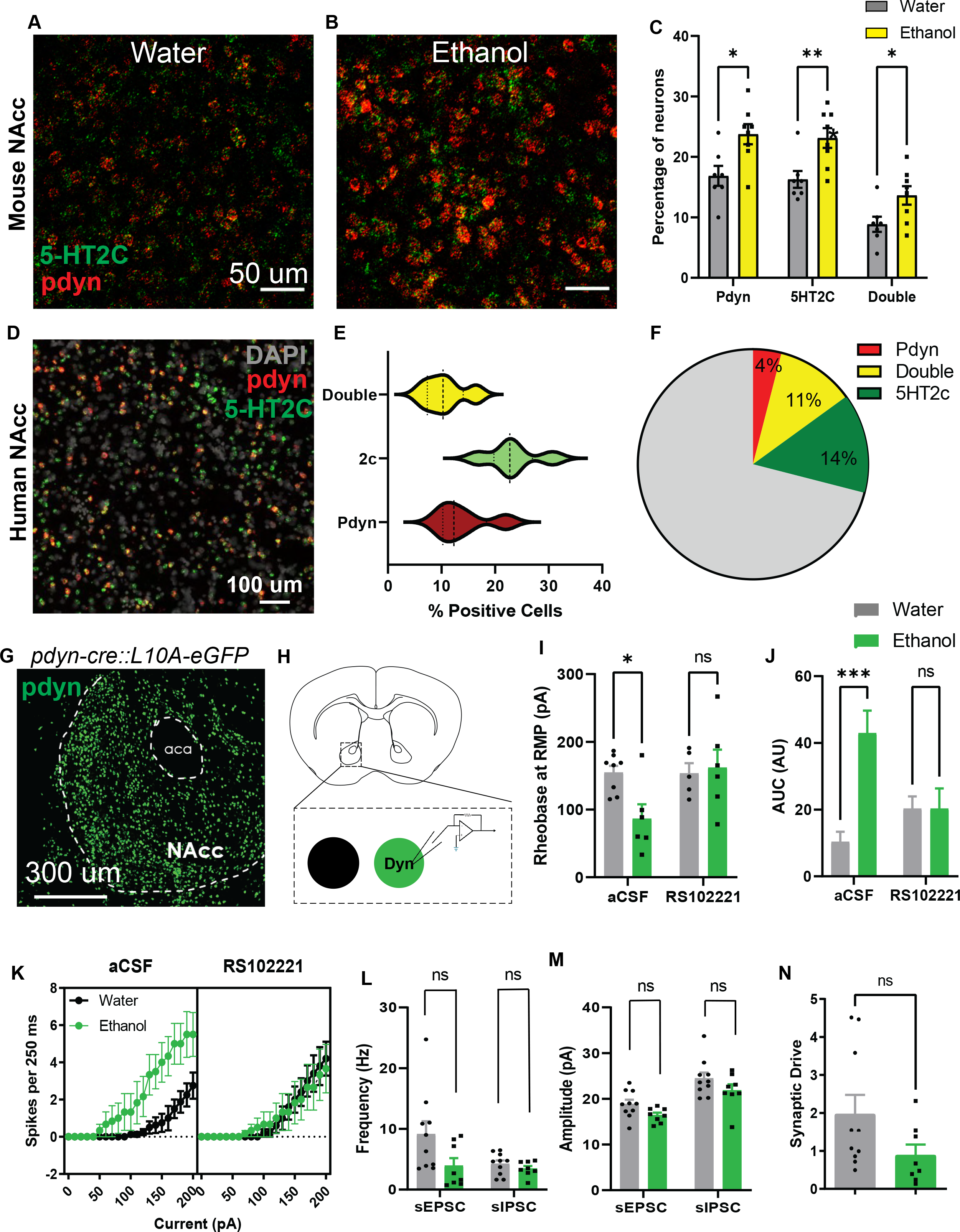
Chronic alcohol upregulates 5-HT2C receptor signaling in NAcc dynorphin neurons. (A-B) Fluorescence in situ hybridization (FISH) images of prodynorphin and 5-HT2C receptors in the NAcc from control and CIE mice (scale bar = 50 µm). (C) Histogram of the percentages of prodynorphin, 5-HT2CR, and double positive neurons in the NAcc in control and CIE mice. (D) Confocal images of neurons expressing 5- HT2C and prodynorphin transcripts captured by FISH in the human NAcc (scalebar = 100 µm). (E) Violin plot depicting the percentages of neurons that express prodynorphin, 5-HT2C, or both in the human NAcc. (F) Pie chart depicting the percentages of neurons that express prodynorphin only, 5-HT2C only, or both in the human NAcc. (G) Confocal image of a brain section of the NAcc taken from a *pdyn-cre::L10A-eGFP* reporter mouse (scale bar = 300 µm). (H) Experimental schematic depicting *ex vivo* electrophysiological recordings in NAcc dynorphin neurons. (I) Histograms of rheobase, (J-K) action potential frequency vs current plots and area under the curve (AUC; AU: arbitrary unit), (L-M) frequency and amplitude of spontaneous EPSCs (sEPSCs) and IPSCs (sIPSCs), and (N) synaptic drive in NAcc dynorphin neurons of control and CIE mice. * p<0.05, ** p<0.01, *** p<0.001, ns: non-significant.

### 5-HTDRN➔NAcc neurons have collateral projections to other cortical and subcortical regions

Our data suggest that inhibiting 5-HT^DRN➔NAcc^ neurons rescued social deficits after CIE, but these neurons may have collateral projections to other regions that drive social behaviors. In order to address this question, we injected *Tph2-p2A-flpO* mice with AAVrg-fDIO-Cre in the NAcc and AAV8.2-hef1α-DIO-synaptophysin-eYFP in the DRN to label axonal processes in downstream regions that may represent collateral projections of 5-HT^DRN➔NAcc^ neurons. We were particularly interested in regions where we had observed elevated c-fos expression after chemogenetic stimulation of 5-HT^DRN^ neurons, including the central lateral nucleus of the thalamus (CL thalamus), NAcc, ACC (layer 1), primary auditory cortex (layer 6a) and BNST. Of those regions, we detected eYFP-labeled processes in the NAcc, BNST, and CL thalamus. The BNST is of particular interest because there is elevated 5-HT_2C_ receptor expression after CIE^11^, although a recent study found that BNST 5-HT_2C_ receptors did not mediate behavioral dysregulation following binge alcohol consumption^18^. We also examined regions outside of this network and found eYFP-labeled axons in the ventral pallidum, ventral tegmental area (VTA), hypothalamus, amygdala, insular cortex, entorhinal cortex, and medial prefrontal cortex. (Extended Data Figure 5). However, since these regions did not exhibit altered c-fos immunoreactivity following chemogenetic stimulation of 5-HT^DRN^ neurons, it is unclear whether they are involved in behavioral deficits following CIE.

**Figure 5:**
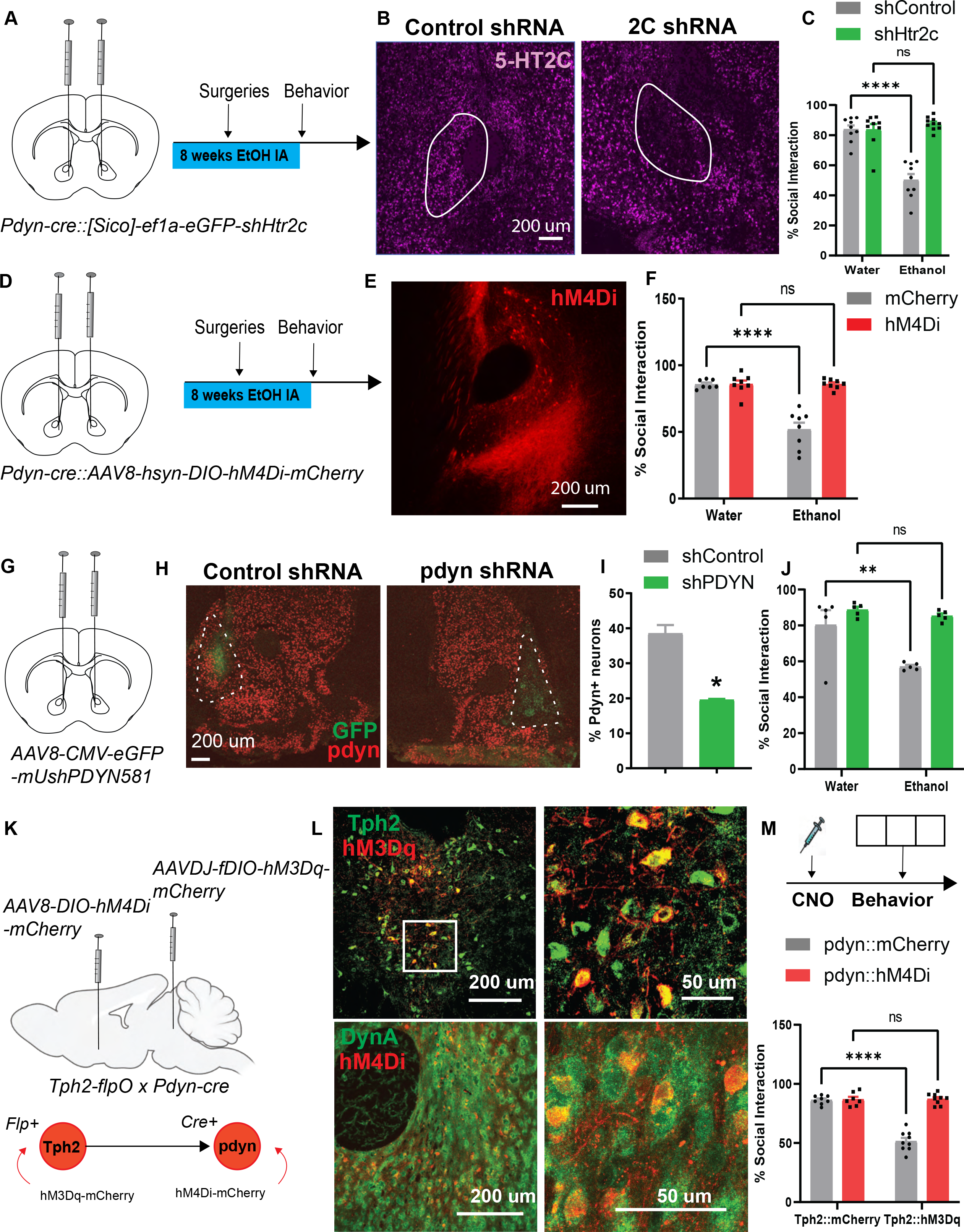
Social deficits in ethanol withdrawal are mediated by activation of 5-HT2C-expressing dynorphin neurons in the NAcc. (A) Viral induction of shRNA directed against the 5-HT2C receptor (*Htr2c*) in NAcc dynorphin neurons and experimental timeline of CIE, surgeries, and behavioral tests. (B) Confocal images of 5-HT2C and prodynorphin FISH in the NAcc depicting 5-HT2C receptor knockdown in dynorphin neurons. (C) Percent time in social interaction showing that 5-HT2C knockdown alleviates social deficits after CIE. (D) Stereotaxic infusion of AAV encoding Cre-inducible Gi-coupled DREADDs in the NAcc of pdyn-cre mice and experimental timeline of CIE, surgeries, and behavioral tests. (E) Confocal image of hM4Di expression in dynorphin neurons in the NAcc (scale bar = 200 µm). (F) Percent time in social interaction indicating that chemogenetic inhibition of NAcc dynorphin neurons alleviates CIE-induced social deficits. (G) Stereotaxic infusion of AAV encoding *pdyn* or scrambled shRNA into the NAcc of C57BL/6J mice exposed to CIE. (H) Validation of *pdyn* knockdown by FISH (scale bar = 200 µm). (I) Histogram of % prodynorphin-expressing neurons in the NAcc in shControl- and shPDYN-injected mice. (J) Percent time in social interaction showing that *pdyn* knockdown rescued social deficits after CIE. (K) Stereotaxic injections into Tph2-flpO::pdyn-cre mice showing viral targeting of hM3Dq to 5-HT^DRN^ neurons and hM4Di to pdyn^NAcc^ neurons. (L) Confocal images showing DREADD expression in 5- HT^DRN^ (top) and pdyn^NAcc^ neurons (bottom) (Left scale bars: 200 µm; Right scale bars: 50 µm) (M) Experimental timeline and percent time in social interaction after CNO. * p<0.05, ** p<0.01, **** p<0.0001, ns: non- significant. *Parts of this figure were made using Biorender*.

### Serotonin release is enhanced in the NAcc during social interaction

To further examine the role of the 5-HT^DRN➔NAcc^ pathway in social deficits after CIE, we next asked whether 5-HT is released in the NAcc during social interaction in ethanol-naïve mice. C57BL/6J mice were stereotaxically injected with a genetically encoded 5-HT biosensor (AAV9- hsyn-5-HT3.5 or GRAB_5-HT3.5_) and implanted with optical fibers in the NAcc for fiber photometry recordings. We observed a significant increase in area under the curve (AUC) in the peri-event plot during the first social approach relative to empty cage investigation (t_7.82_=3.45, p<0.01 with Welch’s correction), suggesting that enhanced 5-HT release in the NAcc coincides with the onset of social interaction. Enhanced activity in 5-HT^DRN➔NAcc^ neurons after CIE may increase basal 5-HT activity in the NAcc and ultimately dysregulate phasic 5-HT release during social interaction, resulting in social deficits (Extended Data Figure 6).

**Figure 6:**
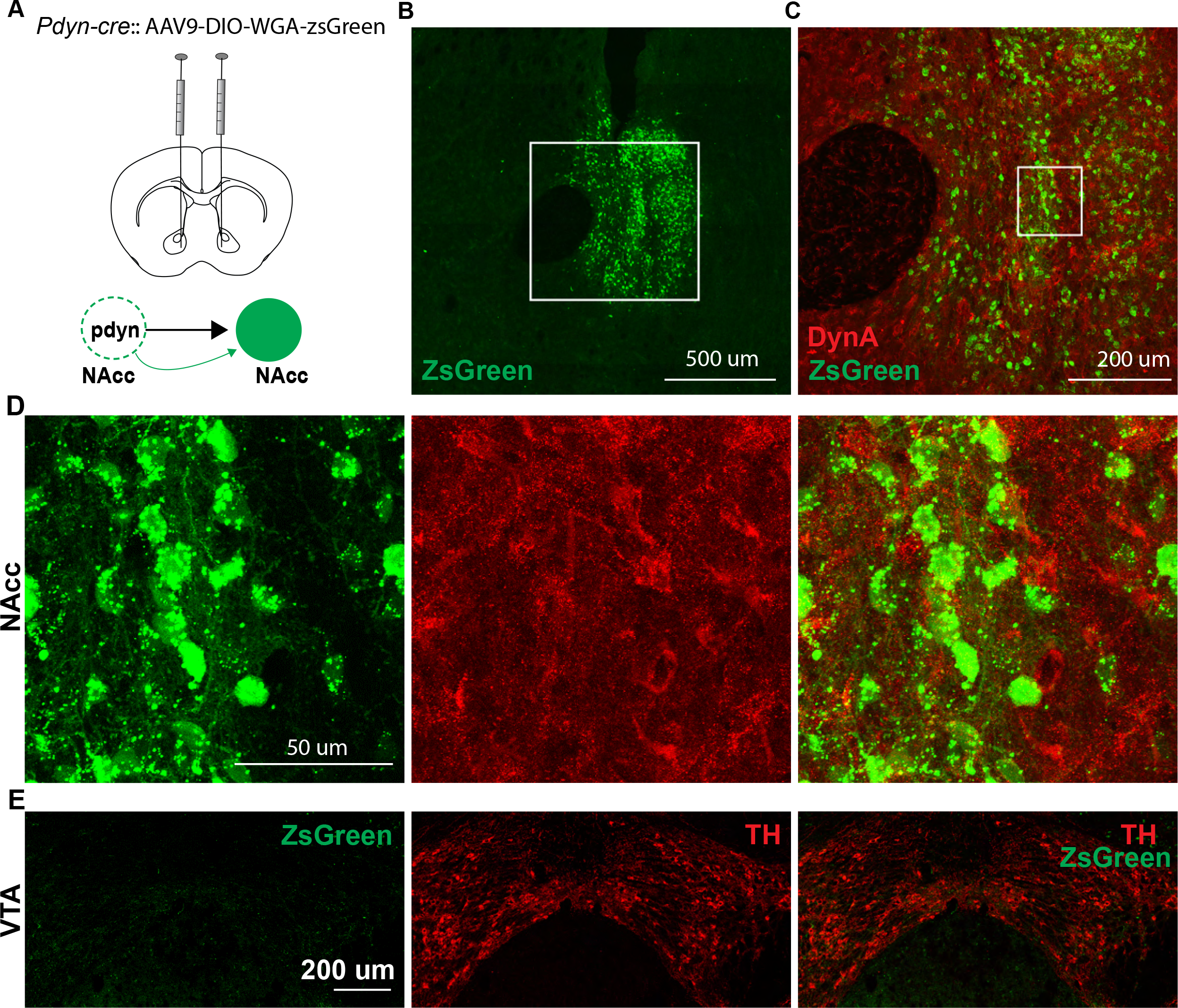
Dynorphin neurons in the NAcc have local synaptic connections. (A) Stereotaxic infusion of AAV expressing Cre-inducible anterograde transsynaptic tracer WGA-ZsGreen in the NAcc of *pdyn-cre* mice. (B) Confocal image of ZsGreen-expressing post-synaptic neurons (scale bar = 500 μm) and (C) colocalization of post-synaptic neurons with dynorphin A in the NAcc (scale bar = 200 μm). (D) Higher magnification (40X) images showing ZsGreen (left), Dynorphin A (center), and overlay (right) in the NAcc (scalebar = 50 μm). (E) Confocal images of ZsGreen, tyrosine hydroxylase (TH) and overlay in the VTA (scale bar = 200 μm).

### 5-HT neurons in the DRN form synaptic connections with NAcc dynorphin neurons and increase neural activity via 5-HT2C receptors

We next injected *Sert-cre* mice with AAV8.2-ef1α-DIO-synaptophysin-eYFP and found that serotonergic fibers originating in the DRN were robustly expressed in the ventral NAcc in close proximity to dynorphin neurons (Figure 3A-B). We also observed synaptic junctions between eYFP-labeled presynaptic terminals and dynorphin A-labeled post-synaptic terminals in the ventral NAcc using transmission electron microscopy (TEM) (Figure 3C). To confirm that ventral NAcc dynorphin neurons receive direct synaptic input from 5-HT^DRN^ neurons, we injected *Sert-cre* mice with the anterograde transsynaptic tracer AAV9-CAG-WGA-ZsGreen in the DRN^19^. This resulted in robust labeling of dynorphin neurons in the ventral NAcc (Figure 3D-F).

A previous study reported that 5-HT_2C_ receptors were enriched in dynorphin-positive cell clusters in the striatum^20^, so we asked whether CIE upregulated 5-HT_2C_ receptors within NAcc dynorphin neurons. Fluorescence *in situ* hybridization (FISH) analysis revealed that CIE increased the percentage of neurons that were double-positive for prodynorphin and 5-HT_2C_ (Figure 4A-C). In contrast, other 5-HT receptors including the 5-HT_1B_ or 5-HT_2A_ were not significantly modified after CIE (Extended Data Figure 7). Striatal 5-HT_2C_ receptors have been previously implicated in anxiety and dysphoria and may promote escalated drinking in ethanol-dependent mice^21^, but a direct link to human AUD has not yet been established. We then asked whether 5-HT_2C_ receptors colocalize with dynorphin neurons in human NAcc sections obtained from the University of Iowa Neurobank (demographic details in Table 1). Overall, 23.25 ± 2.2% of NAcc neurons were 5-HT_2C_-positive, 13.92 ± 2.2% were prodynorphin-positive, and 10.61 ± 1.71% were double positive for 5-HT_2C_ and prodynorphin (Figure 4D-F). These data suggest that the distribution of 5-HT_2C_ receptors in the human NAcc is similar to the mouse, and these receptors may play similar roles in behavioral regulation.

**Figure 7:**
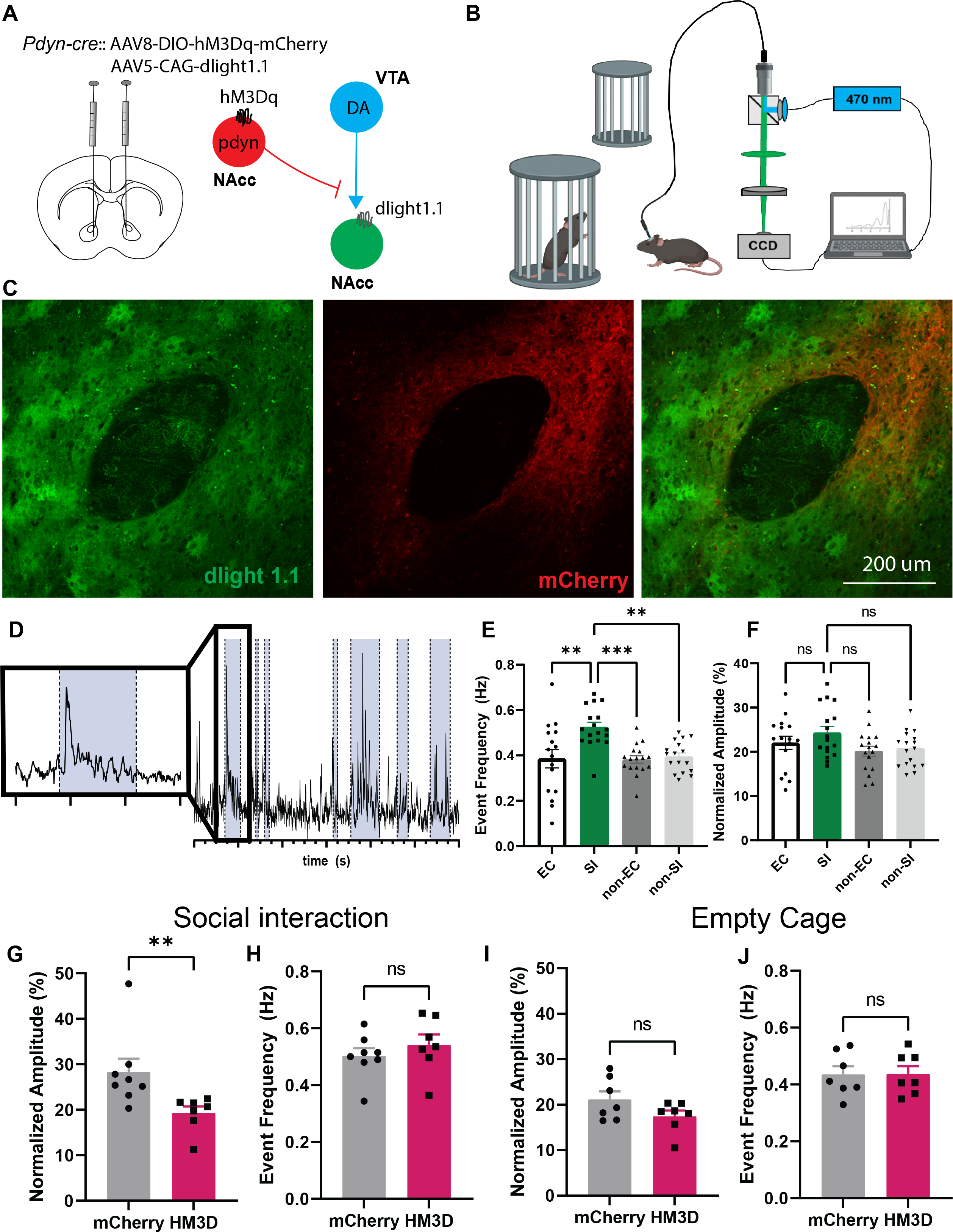
Dynorphin neurons in the NAcc inhibit local dopamine release during social interaction. (A) Stereotaxic infusions of AAV expressing dlight1.1 and Cre-inducible hM3Dq-mCherry into the NAcc of *pdyn-cre* mice. (B) Experimental schematic of fiber photometry recordings during social interaction. (C) Confocal images of dlight1.1 and hM3Dq expression in the NAcc (scale bar = 200 µm). (D) Representative trace of dopamine release events during social interaction, investigation of an empty cage or baseline behavior. Social interactions are highlighted in blue. Time scale: 5 s between two adjacent minor ticks. (E) Histogram of event frequency of dopamine transients during social interaction (SI), empty cage investigation (EC), and exploration of the arena (non-EC: before introduction of the novel mice; non-SI: during the social phase) at baseline (without CNO). (F) Histogram of min/max normalization of mean amplitude of dopamine transients during SI, EC, non-SI and non-EC at baseline. (G-H) Histograms comparing normalized peak amplitude and event frequency during SI in mCherry and hM3Dq groups. (I-J) Histograms comparing peak amplitude and event frequency during EC in mCherry and hM3Dq groups. *p<0.05, ** p<0.01, *** p<0.001, ns: non-significant. *Parts of this figure were made using Biorender*.

**Table 1:** Demographic information for human post-mortem tissue

| **ID Number** | **Age** | **Sex** | **Diagnosis** |
| --- | --- | --- | --- |
| **P3** | 72 | M | Depressive disorder NOS<br>Anxiety disorder NOS |
| **P6** | 60 | F | Major depressive disorder, PTSD, ADD, bulimia |
| **P217** | 74 | M | Depressive disorder NOS |
| **P248** | 67 | F | Depressive disorder NOS |
| **P257** | 69 | F | Generalized Anxiety disorder |

Given that the 5-HT_2C_ receptor is a G-protein-coupled receptor (GPCR) that activates G_q/11_ signaling and promotes membrane depolarization, we expect this upregulation of 5-HT_2C_ receptors to enhance excitability of NAcc dynorphin neurons. As such, we crossed *Pdyn-cre* mice to *L10AeGFP^flox/flox^* mice to use as reporters for *ex vivo* slice electrophysiology experiments (Figure 4G-H). Dynorphin neurons from CIE mice had lower action potential thresholds and enhanced current-induced spiking relative to controls, and this effect was abolished by pre-application of the 5-HT_2C_ receptor antagonist RS102221 (Rheobase: F_1,9_=10.78, p<0.01, Bonferroni t_21_=2.79, p<0.05; Current-induced spiking: F_20,420_ = 2.80, p<0.0001). We also calculated the area under the curve (AUC) of the action potential frequency vs. applied current plot and found AUC was increased in CIE mice but normalized by RS102221 pretreatment (F_1,21_=10.44, p<0.01, Bonferroni t_21_=4.86, p<0.001) (Figure 4I-K). There were no group differences in spontaneous synaptic transmission (inhibitory or excitatory) or synaptic drive (Figure 4L-N), suggesting that ethanol exposure primarily affects the intrinsic excitability of NAcc dynorphin neurons in a 5-HT_2C_ receptor-dependent manner.

### CIE promotes social deficits via 5-HT_2C_ receptors and prodynorphin in the NAcc

We next asked whether genetic ablation of 5-HT_2C_ receptors in NAcc dynorphin neurons could mitigate social deficits in CIE mice. *Pdyn-cre* mice were stereotaxically injected with AAV8-Sico- ef1α-eGFP-Htr2c[shRNA#1] in the NAcc, which expressed eGFP in the absence of Cre and a short-hairpin RNA (shRNA) directed against 5-HT_2C_ receptors in the presence of Cre recombinase. A control scrambled shRNA was also injected into the NAcc of water and CIE mice (shRNA sequence details in Table 2). This resulted in genetic knockdown of 5-HT_2C_ receptors exclusively in dynorphin neurons in the NAcc and alleviated social deficits in CIE mice (F_1,33_=38.07, p<0.0001; Bonferroni post-tests: Water-shControl vs. CIE-shControl, t_33_=7.70, p<0.0001; Water-shHtr2c vs. CIE-shHtr2c, t_33_=0.94, ns) (Figure 5A-C). Chemogenetic inhibition of these NAcc dynorphin neurons produced similar effects (Interaction: F_1,27_=30.62, p<0.0001; Bonferroni post-tests: Water-mCherry vs. CIE-mCherry, t_27_=7.71, p<0.0001; Water-hM4Di vs.

**Table 2:**
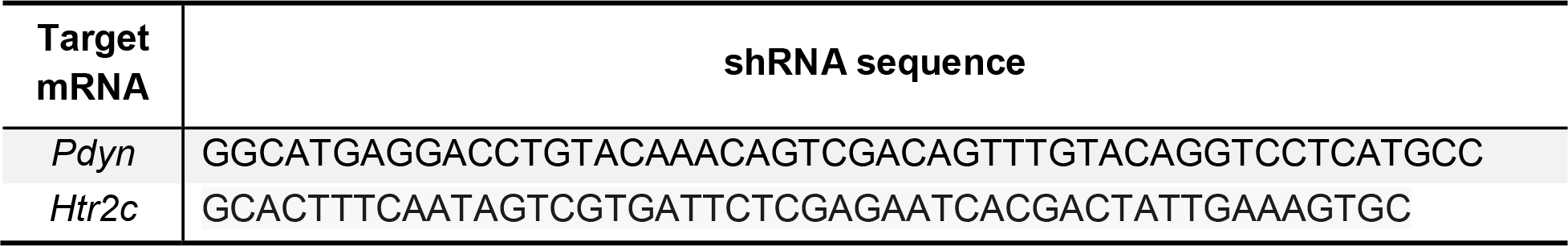
shRNA sequences used in the knockdown experiments

CIE-hM4Di, t_27_=0.01, ns) (Figure 5D-F), suggesting that 5-HT may articulate its aversive effects on social behavior by activating NAcc dynorphin neurons. We then wanted to confirm that dynorphin peptide was mediating these effects since dynorphin-expressing neurons in other brain regions co-express other neuropeptides that are abundant in the striatum including corticotropin-releasing factor (CRF)^22–25^. To do this, we knocked down dynorphin expression in the NAcc with shRNA directed against prodynorphin. AAV8-CMV-shPDYN or scrambled shRNA was stereotaxically injected into the NAcc of C57BL/6J mice exposed to CIE (shRNA sequence details in Table 2). We found that pdyn knockdown in the NAcc normalized time spent in social interaction in CIE mice (Interaction F_1,16_=5.51, p<0.05; Bonferroni post-tests: Water-shControl vs. CIE-shControl, t_16_=3.88, p<0.01; Water-shPDYN vs. CIE-shPDYN, t_16_=0.56, ns) (Figure 5G- J), confirming that dynorphin peptide mediates social deficits after CIE.

### Chemogenetic inhibition of Pdyn^NAcc^ neurons abolishes social deficits induced by 5- HT^DRN^ neuronal stimulation

We have shown so far that stimulation of 5-HT^DRN➔NAcc^ neurons and pdyn^NAcc^ neurons can independently drive social deficits, but it is unclear whether coordinated activity across these two systems is required for the inhibition of social behavior. To test this, we generated a genetic cross between *Tph2-p2A-flpO* and *Pdyn-cre* mice. *Tph2-p2A-flpO*::*Pdyn-cre* mice were stereotaxically injected with AAVDJ-fDIO-hM3Dq-mCherry in the DRN and AAV8-DIO-hM4Di- mCherry in the NAcc. Control mice received AAVDJ-fDIO-mCherry in the DRN and/or AAV8- DIO-mCherry in the NAcc. Here we found that chemogenetic inhibition of ventral NAcc dynorphin neurons mitigated social deficits elicited by activating 5-HT^DRN^ neurons (Interaction: F_1,29_=71.95, p<0.0001, Bonferroni post-tests: Tph2-mCherry::pdyn-mCherry vs. Tph2- hM3Dq::pdyn-mCherry, t_29_=12.11, p<0.0001; Tph2-mCherry::pdyn-hM4Di vs. Tph2- hM3Dq::pdyn-hM4Di, t_29_=0.10, ns) (Figure 5K-M).

### Pdyn^NAcc^ neurons form local circuits and inhibit dopamine release during social interaction

Dynorphin has been previously shown to inhibit dopamine release in the NAcc via presynaptic kappa opioid receptors^26^, but the precise origin of these dynorphinergic inputs is unknown. In another study, optogenetic stimulation of dynorphin in the ventral NAcc induced aversive behavior, but it was suggested that this may be mediated by long-range projections to VTA dopamine neurons^27^. Dynorphin neurons in the NAcc may also inhibit local dopamine release, which would provide a mechanistic framework for understanding their role in social avoidance during ethanol withdrawal. First, we asked whether NAcc dynorphin neurons form local synaptic connections in the NAcc itself by injecting *Pdyn-cre* mice with the anterograde transsynaptic tracer AAV9-CAG-DIO-WGA-ZsGreen to map their post-synaptic targets. A sizable population of non-dynorphin neurons labeled with ZsGreen was detected in the NAcc, suggesting the existence of a local NAcc dynorphinergic circuit (Figure 6A-D). Surprisingly, we did not detect any ZsGreen-labeled cells in the vicinity of dopamine neurons in the VTA, so Pdyn^NAcc^ inputs to VTA dopamine neurons may not be as robust as previously thought (Figure 6E).

We then asked whether chemogenetic stimulation of pdyn^NAcc^ neurons could inhibit local dopamine release in the NAcc in mice actively engaged in social interaction. *Pdyn-cre* mice were co-injected with AAV8-hsyn-DIO-hM3Dq-mcherry and AAV1-CAG-dlight1.1 in the NAcc so that we could simultaneously activate local dynorphin neurons with CNO and detect dopamine release with fiber photometry (Figure 7A-C). The frequency of dopamine transients increased when the animal was actively engaged in social interaction (SI) relative to exploration of empty cage (EC) or when exploring other aspects of the environment (F_3,63_=7.56, p<0.001; Bonferroni post-tests: SI vs EC: t_63_=3.88, p<0.01; SI vs non-EC: t_63_=4.04, p<0.001; SI vs non-SI: t_63_=3.68, p<0.01), but the peak amplitude was unaffected (Figure 7D-F). In a separate session, the mice received intraperitoneal injections of CNO to activate NAcc dynorphin neurons while the mice were engaged in social interaction. Here we found that chemogenetic activation of NAcc dynorphin neurons reduced the peak amplitude of dopamine transients during social interaction (Mann-Whitney U=4, p<0.01) (Figure 7G) but not during exposure to the empty cage (Mann-Whitney U=16, ns) (Figure 7I). On the other hand, there was no effect of chemogenetic manipulation on the frequency of dopamine transients during social interaction (t_13_=0.87, ns) or exploration of the empty cage (t_12_=0.05, ns). The increase in frequency of dopamine transients during social interaction may reflect an increase in burst firing of dopamine neurons in the VTA that project to the NAcc^28^, which is not affected by local dynorphin activity. However, the ability of NAcc dynorphin neurons to reduce the amplitude of these dopamine transients suggests that dynorphin may be acting locally to inhibit release from dopamine terminals. This is consistent with previous microdialysis studies indicating that kappa opioid receptor agonists into the NAcc can reduce extracellular dopamine levels^26^.

## Discussion

Selective serotonin reuptake inhibitors (SSRIs) are commonly prescribed drugs for treating depression and anxiety which frequently present in individuals with AUD^29^. While these drugs are generally thought to mitigate depressive symptoms in these individuals, they can also increase the risk of alcohol relapse during periods of abstinence^30–32^. Although the neural mechanism behind this observation is not abundantly clear, there is some consensus around the idea that alcohol exposure increases serotonergic activity in the DRN and alters neural responses to serotonin release in downstream circuits ^6, 8, 11, 18, 21, 33–36^. Here we demonstrate that ethanol-induced social deficits in males are orchestrated by DRN serotonergic neurons that project to the NAcc and activate 5-HT_2C_ receptors, which are upregulated in dynorphin neurons following CIE. These Gq-coupled 5-HT_2C_ receptors drive excitatory, postsynaptic signaling and potentiates dynorphin neuronal activity, which in turn inhibits dopamine release during social interaction. Thus serotonin, acting via the 5-HT_2C_-dynorphin circuit, has an inhibitory effect on dopamine release in the NAcc, which may reduce the motivation to engage in social interaction. Interestingly, both serotonin and dopamine release increased during social interaction in ethanol-naïve animals. This suggests that serotonin release during social interaction may act as a molecular brake on dopamine release that eventually turns off this signal via the 5-HT_2C_ receptor as the social encounter becomes less novel and more familiar. After chronic ethanol, however, this 5-HT^DRN^➔pdyn^NAcc^ circuit goes into overdrive, which impairs dopamine release in the NAcc and degrades the reward value of social interaction.

5-HT^DRN➔NAcc^ neurons also have collateral projections to other regions that could potentially drive social behaviors, including the BNST which is enriched in 5-HT_2C_ receptors that are upregulated in mice following CIE^11^. However, our RNAi data suggests that CIE-induced social deficits are primarily driven by 5-HT_2C_ receptors in NAcc dynorphin neurons. Interestingly, it has also been reported that 5-HT inputs to the BNST promote fear and anxiety-like behaviors in ethanol-naïve mice^37^, so elevated 5-HT_2C_ signaling in the BNST could mediate some other behavioral endpoints associated with excessive alcohol use.

The idea that excessive serotonergic drive in the NAcc inhibits social behavior seems to contradict some of the previous literature in which 5-HT inputs to the NAcc alleviate social deficits in mouse models of autism^15^. However, it is important to note that 5-HT can have divergent effects depending on the distribution and density of specific 5-HT receptor subtypes, which can be pre- or post-synaptic and activate different G protein-coupled signaling pathways. In 16p11.2 mice, the socially rewarding effects of 5-HT^DRN^ stimulation were mediated by 5-HT_1B_ receptors, which are presynaptic Gi-coupled receptors that inhibit release of other neurotransmitters. In the context of CIE, there is upregulation of 5-HT_2C_ receptors in NAcc dynorphin neurons which tends to increase dynorphinergic output and cause a net inhibitory effect on phasic dopamine release during social behavior. While 5-HT signaling in the NAcc may promote social reward when 5-HT_1B_ receptors are abundant, upregulation of 5-HT_2C_ receptors after chronic alcohol may convert 5-HT to an aversive signal.

In total, these studies demonstrate that 5-HT neurons that project to the NAcc can modulate social behavior by activating 5-HT_2C_ receptors in dynorphin neurons. Chronic alcohol engages this 5-HT^DRN^➔Dyn^NAcc^ circuit by enhancing activity in 5-HT^DRN^ neurons and increasing 5-HT_2C_ receptor signaling in NAcc dynorphin neurons, which ultimately suppresses dopamine release during social interaction and promotes social deficits (Figure 8). Overall, the results of this study strongly suggest that SSRIs should be used with caution in individuals with AUD, especially in those that are recently abstinent from alcohol as these individuals may experience more negative side effects from taking antidepressants that could ultimately lead to relapse.

**Figure 8:**
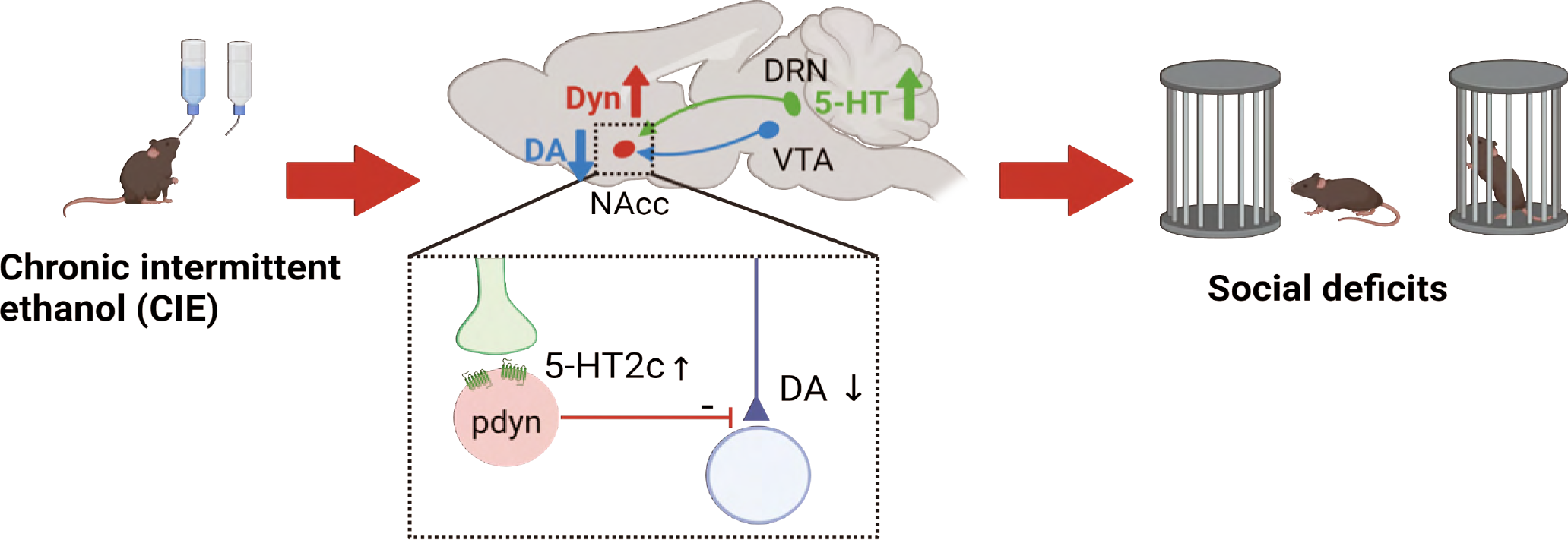
Conceptual model depicting how chronic alcohol use can induce social deficits. Chronic ethanol exposure drives hyperactivity in 5-HT DRN neurons, which activate 5-HT2C receptors in dynorphin neurons in the NAcc. This in turn inhibits dopamine release from dopamine terminals in the NAcc during social investigation, which ultimately inhibits that behavior. *Parts of this figure were made using Biorender.*

Future studies would be warranted to determine whether SSRIs could be more effective in individuals with AUD when combined with short-term use of a kappa opioid or 5-HT_2C_ receptor antagonist.

## Methods

### 2.1 Animals

In these experiments, we used adult male and female C57BL/6J mice from Jackson Labs (Bar Harbor, ME, USA), as well as transgenic *Sert-cre* (Jackson Labs, 014554), *Pdyn-ires-cre* (Jackson Labs, 027958), and *EGFP-L10a* mice (Jackson Labs, 024750) which were bred in-house. Tph2-p2A-flpO mice were developed at the University of Iowa Genome Editing Core Facility. All mice were maintained at a temperature- and humidity-controlled AAALAC approved vivarium at the University of Iowa College of Medicine with *ad libitum* access to food and water and 24 hour on-call veterinary care.

#### Generation of Tph2-p2A-flpO mice

To be able to target subpopulations of 5-HT neurons, we generated a *Tph2-p2A-flpO* mouse line that expresses flpO recombinase from the endogenous *Tph2* locus^38^. CRISPR-Cas9 endonuclease-mediated genome editing was used to insert a p2A-flpO-recombinase sequence 3’ in coding exon 11 of the *Tph2* locus (Figure 2I). The self-cleaving p2A sequence allows for expression of the endogenous Tph2 protein and flpO recombinase. B6SJLF1/J mice (Jackson Labs, 100012) were used in the production of *Tph2-p2A-flpO* mice. To validate this mouse line, we performed fluorescence *in situ* hybridization (FISH) and immunofluorescence assays (Extended Data Figure 4). In FISH, RNAscope probes Mm-Tph2-C2 and Flp from ACDBio (Newark, CA, USA; Table 3) were used to assess colocalization of Tph2 with flpO RNA in the DRN. In immunohistochemistry, we first stereotaxically injected Tph2-p2A-flpO mice with AAV5- ef1α-fDIO-mCherry in the DRN, and 4 weeks later, we examined colocalization of Tph2 with mCherry (see antibody details in Table 4).

**Table 3:** RNAscope probes used in fluorescence *in situ* hybridization

| **Species** | **mRNA** | **ACDBio Cat No.** |
| --- | --- | --- |
| Mouse | *Pdyn* | 318771 |
|  | *Htr2c* | 401001-C3 |
|  | *Htr2a* | 401291-C3 |
|  | *Htr1b* | 315861 |
|  | *Flp (customized)* | 448191 |
|  | *Tph2* | 318691-C2 |
| Human | *Pdyn* | 507161-C2 |
|  | *Htr2c* | 450651 |

**Table 4:**
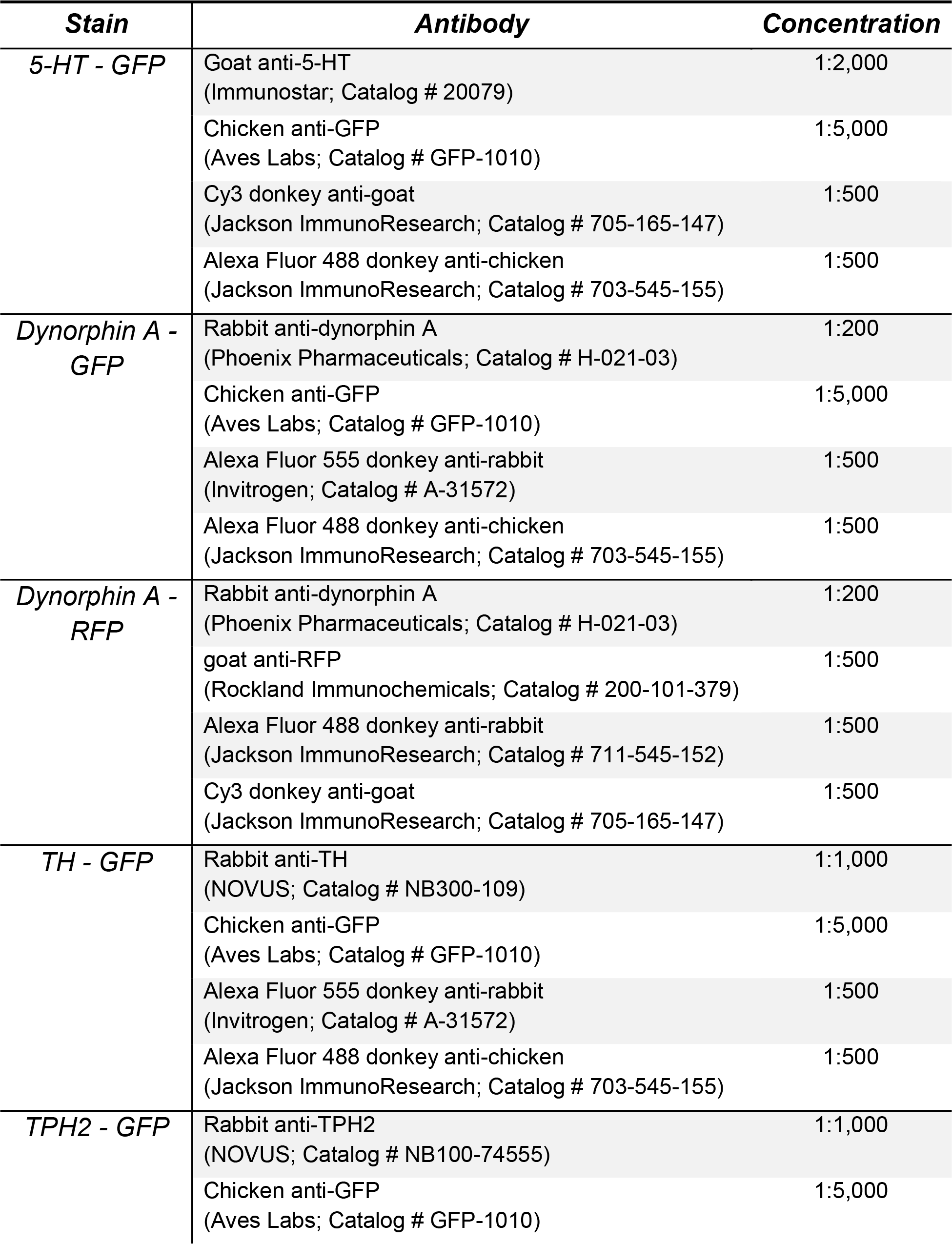
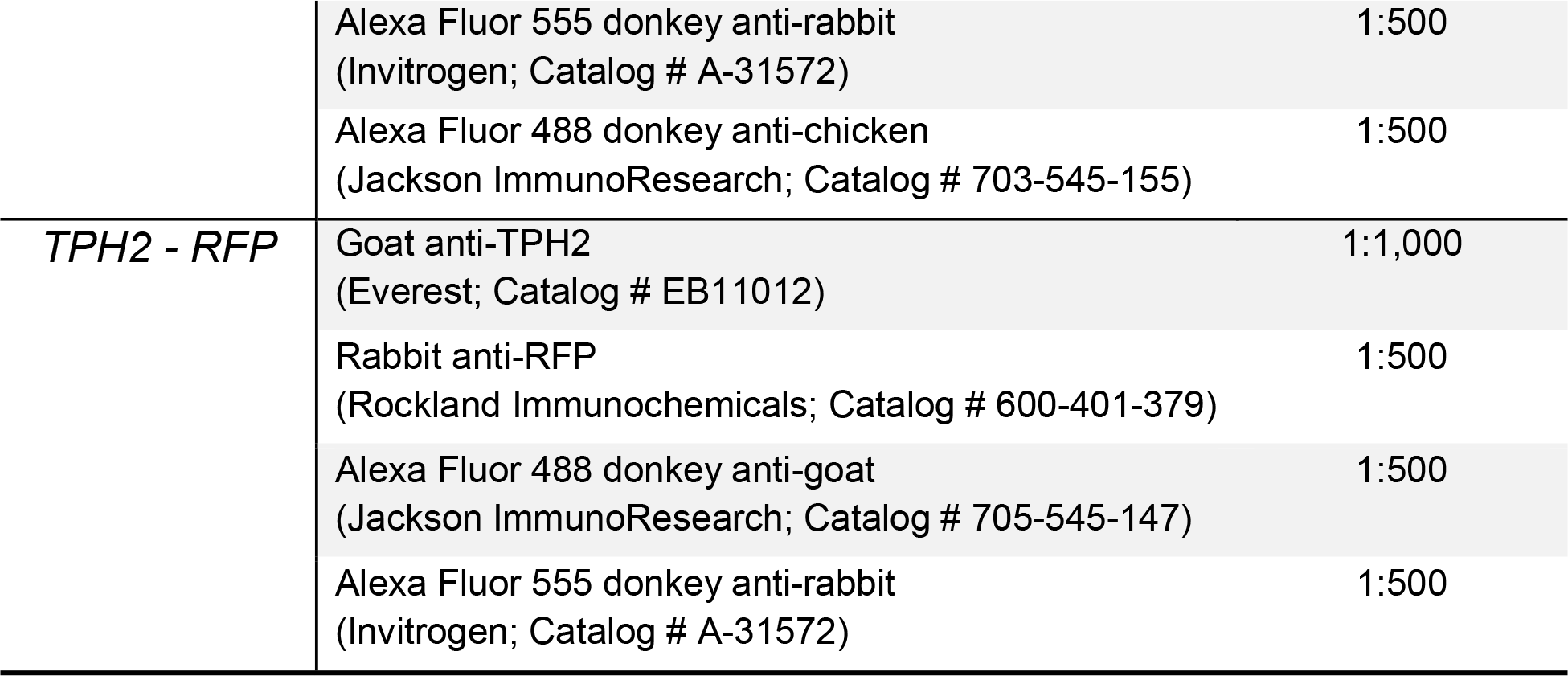
Antibodies used in immunofluorescence

| ***Stain*** | ***Antibody*** | ***Concentration*** |
| --- | --- | --- |
| ***5-HT - GFP*** | Goat anti-5-HT<br>(Immunostar; Catalog # 20079) | 1:2,000 |
|  | Chicken anti-GFP<br>(Aves Labs; Catalog # GFP-1010) | 1:5,000 |
|  | Cy3 donkey anti-goat<br>(Jackson ImmunoResearch; Catalog # 705-165-147) | 1:500 |
|  | Alexa Fluor 488 donkey anti-chicken<br>(Jackson ImmunoResearch; Catalog # 703-545-155) | 1:500 |
| ***Dynorphin A - GFP*** | Rabbit anti-dynorphin A<br>(Phoenix Pharmaceuticals; Catalog # H-021-03) | 1:200 |
|  | Chicken anti-GFP<br>(Aves Labs; Catalog # GFP-1010) | 1:5,000 |
|  | Alexa Fluor 555 donkey anti-rabbit<br>(Invitrogen; Catalog # A-31572) | 1:500 |
|  | Alexa Fluor 488 donkey anti-chicken<br>(Jackson ImmunoResearch; Catalog # 703-545-155) | 1:500 |
| ***Dynorphin A - RFP*** | Rabbit anti-dynorphin A<br>(Phoenix Pharmaceuticals; Catalog # H-021-03) | 1:200 |
|  | goat anti-RFP<br>(Rockland Immunochemicals; Catalog # 200-101-379) | 1:500 |
|  | Alexa Fluor 488 donkey anti-rabbit<br>(Jackson ImmunoResearch; Catalog # 711-545-152) | 1:500 |
|  | Cy3 donkey anti-goat<br>(Jackson ImmunoResearch; Catalog # 705-165-147) | 1:500 |
| ***TH - GFP*** | Rabbit anti-TH<br>(NOVUS; Catalog # NB300-109) | 1:1,000 |
|  | Chicken anti-GFP<br>(Aves Labs; Catalog # GFP-1010) | 1:5,000 |
|  | Alexa Fluor 555 donkey anti-rabbit<br>(Invitrogen; Catalog # A-31572) | 1:500 |
|  | Alexa Fluor 488 donkey anti-chicken<br>(Jackson ImmunoResearch; Catalog # 703-545-155) | 1:500 |
| ***TPH2 - GFP*** | Rabbit anti-TPH2<br>(NOVUS; Catalog # NB100-74555) | 1:1,000 |
|  | Chicken anti-GFP<br>(Aves Labs; Catalog # GFP-1010) | 1:5,000 |
|  | Alexa Fluor 555 donkey anti-rabbit<br>(Invitrogen; Catalog # A-31572) | 1:500 |
|  | Alexa Fluor 488 donkey anti-chicken<br>(Jackson ImmunoResearch; Catalog # 703-545-155) | 1:500 |
| *TPH2 - RFP* | Goat anti-TPH2<br>(Everest; Catalog # EB11012) | 1:1,000 |
|  | Rabbit anti-RFP<br>(Rockland Immunochemicals; Catalog # 600-401-379) | 1:500 |
|  | Alexa Fluor 488 donkey anti-goat<br>(Jackson ImmunoResearch; Catalog # 705-545-147) | 1:500 |
|  | Alexa Fluor 555 donkey anti-rabbit<br>(Invitrogen; Catalog # A-31572) | 1:500 |

### 2.2 Drugs

#### Chemogenetic experiments

Clozapine-N-oxide dihydrochloride (CNO; Hello Bio, Princeton, NJ, USA) was dissolved in sterile, 0.9% saline to a concentration of 0.30 mg/ml and injected at 10 ml/kg, i.p., 30 min before the beginning of behavioral experiments for a dose of 3 mg/kg. For c-fos studies, CNO was injected 2 hours before perfusion and brain extraction. Deschloroclozapine dihydrochloride (DCZ; Hello Bio) was dissolved in sterile, 0.9% saline to a concentration of 10 µg/ml and injected at 10 ml/kg, i.p., 20 min before the beginning of behavioral experiments for a final dose of 100 µg/kg, which was previously found to induce neural activity in hM3Dq-expressing neurons in mice and monkeys^17^.

#### Electrophysiology and two-photon experiments

RS102221 hydrochloride (Tocris Bioscience, Minneapolis, MN, USA) was first made into a 100 mM stock solution in dimethyl sulfoxide (DMSO), and then diluted to a final concentration of 1 µM in artificial cerebral spinal fluid (aCSF). Serotonin hydrochloride (5-HT; Thermo Fisher Scientific, Waltham, MA, USA) was first made into a 20 mM stock solution in ddH_2_O, and then diluted to a final concentration of 20 µM in aCSF.

### 2.3 Viral constructs

Most of the AAVs used in this study were obtained from Addgene (Watertown, MA, USA), Stanford University Gene Vector and Virus Core (Stanford, CA, USA). AAV2/8-CMV-eGFP- mUsh-Pdyn581 and AAV8-CMV-eGFP-mUsh-Control were generated at the University of Iowa Viral Vector Core using plasmids provided by Brigitte Kieffer at McGill University (now at University of Strasbourg). AAV8-Sico-ef1α-eGFPHtr2c[shRNA#1] and AAV8-Sico-ef1α- eGFPScramble[shRNA#1] were generated and packaged by VectorBuilder (Chicago, IL, USA). AAV8.2-ef1α-DIO-synaptophysin-eYFP was provided by Rachel Neve at the Massachusetts General Hospital (MGH) Gene Technology Core (Boston, MA, USA). AAV-CAG-WGA-ZsGreen plasmid was provided by Qi Wu at Baylor College of Medicine and packaged at the University of Iowa viral vector core. AAV9-hsyn-5-HT3.5 (GRAB_5-HT3.5_) was produced and packaged at WZ Biosciences (Columbia, MD, USA). See Table 5 for details about viral constructs.

**Table 5:** Viral constructs

| AAV | Virus Source | Packaging | Cat No. | Titer |
| --- | --- | --- | --- | --- |
| AAVrg-hsyn-DIO-hM3D(Gq)-mCherry | Addgene | – | 44361-AAVrg | $7 \times 10^{12}$ vg/ml |
| AAVrg-hsyn-DIO-hM4D(Gi)-mCherry | Addgene | – | 44362-AAVrg | $8 \times 10^{12}$ vg/ml |
| AAVrg-hsyn-DIO-mCherry | Addgene | – | 50459-AAVrg | $7 \times 10^{12}$ vg/ml |
| AAV8-hsyn-DIO-hM3D(Gq)-mCherry | Addgene | – | 44361-AAV8 | $4 \times 10^{12}$ vg/ml |
| AAV8-hsyn-DIO-hM4D(Gi)-mCherry | Addgene | – | 44362-AAV8 | $1 \times 10^{13}$ vg/ml |
| AAV8-hsyn-DIO-mCherry | Addgene | – | 50459-AAV8 | $1 \times 10^{13}$ vg/ml |
| AAVDJ-hsyn-fDIO-hM3D(Gq)-mCherry | Stanford Gene Vector and Virus Core | – | GVVC-AAV-153 | $2 \times 10^{13}$ vg/ml |
| AAV5-ef1α-fDIO-mCherry | Addgene | – | 114471-AAV5 | $7 \times 10^{12}$ vg/ml |
| AAVrg-EF1α-fDIO-Cre | Addgene | – | 121675-AAVrg | $7 \times 10^{12}$ vg/ml |
| AAV5-CAG-dlight1.1 | Addgene | – | 111067-AAV5 | $7 \times 10^{12}$ vg/ml |
| AAV2/8-CMV-eGFP-mUsh-Pdyn581 | Brigitte Kieffer at McGill University | U Iowa Viral Vector Core | – | $8.7 \times 10^{12}$ vg/ml |
| AAV2/8-CMV-eGFP-mUsh-Control | Brigitte Kieffer at McGill University | U Iowa Viral Vector Core | – | $6.4 \times 10^{12}$ vg/ml |
| AAV8-Sico-ef1α-eGFP<sup>Htr2c</sup>[shRNA#1] | VectorBuilder | VectorBuilder | – | $2 \times 10^{11}$ gc/ml |
| AAV8-Sico-ef1α-eGFP<sup>Scramble</sup>[shRNA#1] | VectorBuilder | VectorBuilder | – | $2 \times 10^{11}$ gc/ml |
| AAV8.2-ef1α-DIO-synaptophysin-eYFP | Rachel Neve at the Massachusetts General Hospital | – | RN2 | $2.19 \times 10^{13}$ vg/ml |
| AAV2/9-CAG-DIO-WGA-ZsGreen | Qi Wu at Baylor College of Medicine | U Iowa Viral Vector Core | – | $1.12 \times 10^{14}$ vg/ml |
| AAV9-hSyn-5-HT3.5 (GRAB<sub>5-HT3.5</sub>) | WZ Biosciences | – | – | $3.36 \times 10^{13}$ gc/ml |

### 2.4 Stereotaxic surgeries

The mouse was placed in an Angle Two stereotaxic frame (Leica Biosystems, Wetzlar, Germany) and maintained on a heated platform under 5% (v/v) isoflurane anesthesia as described previously^37^. For intracranial injections of AAV, a 1 μl Neuros syringe needle (Model: 7001 KH; Hamilton Company, Reno, NV, USA) was positioned over the brain region of interest (DRN: AP=-4.65, ML= 0.00, DV=-3.30, θ=23.58°; NAcc: AP=0.98, ML=±0.70, DV=-4.55, θ=0°, relative to the Bregma) and the virus was infused with a Stoelting Quintessential Stereotaxic Injector (Wood Dale, IL, USA) at a rate of 100 nl/min (total volume: 800 nL in the DRN; 400 nL/side in the NAcc). The needle remained in place for an additional 10 min to allow for diffusion of the virus before removal. The mice were administered meloxicam (4 mg/kg, s.c.) every 24 hours for 48-72 hours to minimize post-operative pain and discomfort. Mice were allowed at least 7 days to recover before experiments began. For fiber photometry 5-HT biosensor experiments, AAV9-hsyn-5-HT3.5 was first 1:1 diluted with sterile, 0.9% saline, and then the mouse was injected unilaterally with 1 µl total volume of the diluted virus into the right NAcc (AP=0.98, ML=0.70, DV=-4.55, θ=0°). Following viral infusion, a fiber optic cannula (ferrule diameter 2.5 mm, fiber core diameter 400 µM, length 5 mm, NA 0.39, Amuza, San Diego, CA, USA) was placed with the terminal of the fiber at the same coordinates. The cannula was sealed in place with dental cement. For fiber photometry dopamine biosensor experiments, the total volume of 400 nl of a 1:1 mixture of AAV5-CAG-dlight1.1 and AAV8-hsyn-DIO-mCherry or AAV8-hsyn-DIO-hM3Di-mCherry was injected unilaterally into the left NAcc (AP=1.30, ML=- 1.25, DV=-4.25, θ=0°). A fiber optic cannula (ferrule diameter 1.25 mm, fiber core diameter 200 µM, length 5 mm, NA 0.37, Neurophotometrics, San Diego, CA, USA) was then placed and sealed.

### 2.5 Chronic Intermittent access to ethanol (CIE)

Mice were singly housed starting at 7-8 weeks of age in specialty cages with 2 holes at the front of the cage designed to fit two water bottles containing either tap water or 20% w/v ethanol.

Water control mice were given tap water only. Water and ethanol bottles were given between 9:00-10:00 AM on MWF and replaced with two water bottles 24 hours later. The configuration of the bottles (left v right) was switched between drinking sessions to account for side preferences. Two drip bottles for water and ethanol attached to an empty cage were included to measure loss via drip. Water and ethanol consumption over each 24-hour drinking session was calculated as follows:

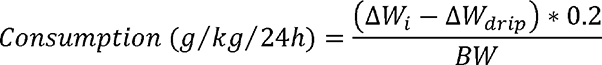

where ΔW_i_ is the weight change of the water or ethanol bottle in grams, ΔW_drip_ is the weight change of the drip bottle, and BW is the body weight of the mouse in kg.

### 2.6 Behavior

#### Social interaction test

The social interaction test was conducted 24 hours after the last ethanol drinking session and 30 min after injection with CNO (in chemogenetic experiments only), which has been described in detail before^11, 12, 34^. Briefly, the test was conducted in a three-chambered Plexiglas container (L × W × H: 64 × 42 × 21 cm) divided by 2 panels with a small opening at the base of each to allow movement between the chambers^39^. Behavior was recorded by an overhead camera with Media Recorder software (Noldus, Leesburg, VA, USA). At the start of each testing session, the test mouse was placed in the center chamber and allowed free movement/exploration throughout the 3 compartments for 10 min. The mouse was then placed back in the center chamber with the doors closed and a holding cage (i.e., a wire mesh pen cup, 11 cm high, open end: 10 cm in diameter; closed end: 8 cm in diameter) was placed in the center of each of the two outermost chambers. A novel conspecific of the same age and sex was placed inside the holding cage on one side while the other cage remained empty. The doors were then re-opened and the test mouse was allowed to roam freely between chambers for 10 more min. The time spent interacting with the stranger mouse and investigating the empty cage was scored by a trained observer blinded to the experimental condition. The percentage of time spent in social interaction or investigating the empty cage during the 10-min session was then calculated for each mouse. The placement of the stranger was alternated between trials to control for side preferences.

#### Elevated plus maze (EPM) test

The Plexiglas elevated plus maze had two open arms (35 x 5 cm each) and two closed arms (35 x 5 cm each, with 20-cm-tall dark side walls), interconnected via a center zone (5 x 5 cm) and elevated by 60 cm from the floor. The test was conducted under dim light (∼20 lux in the open arms and ∼5 lux in the closed arms), starting when the mouse was being placed in the center zone. The mouse was allowed to freely move and explore in the maze for 5 min, and their activity was recorded with a camera mounted above the maze. The videos were then analyzed with Ethovision XT14 software (Noldus), and the following parameters were compared: time in open arms, probability of activity in open arms, latency to open arms, and total distance moved.

#### Open field test

The open field test was conducted in an opaque Plexiglas arena (L x W x H: 50 x 50 x 25 cm), situated in a custom-made sound-attenuating wood box. The arena was placed under dim light (∼20 lux in the center of the arena), with a camera mounted above. The mouse was placed in a corner of the arena and allowed to freely move and explore for 30 min. All the activity was recorded and analyzed with Ethovision XT14 software, by comparing the following parameters: latency to the center, center entries, and total distance moved. The center was defined as the central 15% of the arena.

#### Sucrose preference test

Mice were placed in a PhenoTyper 3000 observation cage (Noldus), consisting of a square chamber (30 x 30 cm) with clear Plexiglas walls, an overhead camera, a feeder with *ad libitum* access to chow, and 2 water bottles on the wall opposite to the feeder, which were both equipped with lickometers that automatically detect contacts made between the mouse and the water spouts. The data from the lickometers and the overhead camera were processed with Ethovision XT14 software. The mouse was allowed to freely explore the chamber for 1 hour on each of 4 training days, with *ad libitum* access to tap water and 5% sucrose solution from the 2 bottles. The placements of the water and sucrose bottles were switched every day to avoid place preferences. The final test lasted for 20 min on the 5^th^ day.

Sucrose preference was calculated as follows:

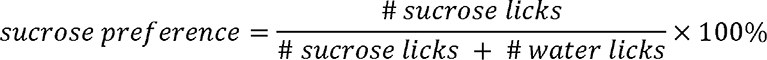

After this 5-day sucrose preference test, the mice were exposed to another cycle of CIE. At 24 hours after the last ethanol exposure (ethanol withdrawal; EW), they were given ad libitum access to 5% sucrose and water for 20 min.

### 2.7 Whole-cell patch clamp electrophysiology

Procedures for whole-cell patch clamp electrophysiology have been described in detail elsewhere^11, 37, 40, 41^. Briefly, mice were decapitated under isoflurane anesthesia 24 h after the last ethanol drinking session. Brains were extracted into a solution of ice-cold sucrose-aCSF (in mM): 194 sucrose, 20 NaCl, 4.4 KCl, 2 CaCl_2_, 1 MgCl_2_, 1.2 NaH_2_PO_4_, 10 glucose, and 26 NaHCO_3_, saturated with 95% O_2_/5% CO_2_, and sectioned at 0.07 mm/s on a Leica 1200S vibratome to obtain 300 μm coronal slices of the DRN or NAcc, which were incubated in a heated holding chamber containing normal, oxygenated aCSF (in mM):124 NaCl, 4.4 KCl, 2 CaCl_2_, 1.2 MgSO_4_, 1 NaH_2_PO_4_, 10.0 glucose, and 26.0 NaHCO_3_, maintained at 30 ± 1 °C for at least 1 h before recording. The slices were then transferred to a recording chamber containing normal, oxygenated aCSF, maintained at 28–30 °C, delivered at a flow rate of 2 ml/min.

Neurons were visualized using infrared differential interference contrast (DIC), video-enhanced microscopy (Olympus, Tokyo, Japan) and their genetic identity as dynorphin or 5-HT neurons was confirmed using GFP and RFP filter cube sets (Olympus). Slices were perfused in normal aCSF in the recording chamber for at least 30 min before whole-cell patch clamp recordings were performed. Borosilicate electrodes were pulled with a Flaming-Brown micropipette puller (Sutter Instruments, Novato, CA, USA) and only those with a pipette resistance between 3 and 6 MΩ were used to patch cells. These pipettes were filled with K-gluconate internal solution (in mM: 135 K-gluconate, 5 NaCl, 2 MgCl_2_, 10 HEPES, 0.6 EGTA, 4 ATP, and 0.4 GTP; pH = 7.35, 290 mOsmol/kg) for excitability experiments. In experiments to examine synaptic transmission, cesium methanesulfonate-based internal solution was used (in mM: 135 cesium methanesulfonate, 10 KCl, 1 MgCl_2_, 10 HEPES, 0.2 EGTA, 4 Mg-ATP, 0.3 Na_2_-GTP, 20 Na_2_- phosphocreatine; pH = 7.30, 290 mOsmol/kg, with 1 mg/ml QX-314).

Signals were acquired using a MultiClamp 700B amplifier and digitized with a Digidata 1550B digitizer (Molecular Devices, San Jose, CA, USA). Signals were then analyzed with pClamp 11 software (Molecular Devices). Excitability experiments were conducted in the current clamp mode with the holding current set to zero. The rheobase was determined using a current ramp protocol to measure the minimum current required to induce firing. The V-I plot was determined by the numbers of spikes generated within 250 ms during a series of discrete 10 pA current steps from 0 to 200 pA. In a subset of experiments, the 5-HT_2C_ receptor antagonist RS102221 (1 μM) was bath applied for 10 min before the rheobase and VI plot experiments were performed. Synaptic transmission was assessed in the voltage clamp mode. Spontaneous excitatory postsynaptic currents (sEPSCs) were recorded continuously for 2 min while the membrane potential was maintained at -55 mV; in the same cell, spontaneous inhibitory postsynaptic currents (sIPSCs) were recorded while the cell was held at 10 mV. Synaptic drive was calculated as follows:

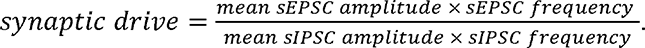

### 2.8 Fluorescence In situ hybridization (FISH)

#### Mouse tissue

Animals were deeply anesthetized with isoflurane before decapitation. Brains were then rapidly isolated, frozen on dry ice, and embedded in tissue freezing media (Tissue-Tek O.C.T. compound, Sakura Finetek USA, Torrance, CA). 16 µm NAcc coronal sections were obtained with a Leica CM3050S cryostat and mounted onto SuperFrost Plus glass slides (VWR, Radnor, PA, USA). The sections were collected in a way that each slide contained rostral, middle, and caudal NAcc sections. Once mounted, slices were rapidly refrozen and stored at −80°C before the FISH procedure. Multiplexed FISH was performed using the ACDBio RNAscope V2 reagents and protocols (Cat No: 323100; Newark, CA, USA). Briefly, slices were fixed in 10% chilled neutral buffered formalin (MilliporeSigma, Burlington, MA, USA) at 4°C for 20 min and dehydrated serially in ethanol (50%, 70%, and 100%). The slides were then treated with RNAscope hydrogen peroxide (10 min) and digested using Protease IV for 30 min at room temperature. After the pretreatment steps, respective mRNA probes for mouse *Pdyn*, *Htr1b*, *Htr2a* and *Htr2c* were applied (see probe details in Table 3). A series of amplification and detection reagents were applied to the tissue sections according to the manufacturer’s instructions. Sections were counterstained using DAPI provided in the detection reagent kits and mounted in ProLong Gold mounting media (P36930, Thermo Fisher Scientific). The images were obtained using an Olympus FV3000 confocal laser scanning microscope with a 40X magnification oil immersion objective. The confocal images were processed in ImageJ Fiji software. Cell detection and RNAscope puncta quantification were performed using a custom pipeline in CellProfiler (version 3.1.5) and QuPath (version 0.4.2).

#### Human tissue

Frozen NAcc brain blocks were obtained from the Iowa Neurobank for FISH analysis (Table 1). 20 µm sections were obtained and mounted on SuperFrost Plus glass slides in the cryostat chamber and the slides were stored at −80°C before the FISH procedure. A similar protocol was used as described above with human-specific probes for Pdyn and Htr2c mRNA (Table 3). The slides were scanned using an Olympus SLIDEVIEW VS200 digital slide scanner and the images were analyzed in QuPath (version 0.4.2), including cell detection and RNAscope puncta quantification.

### 2.9 Immunofluorescence and Confocal Imaging

Deeply anesthetized mice (with tribromoethanol, 250 mg/kg, i.p.) were transcardially perfused using PBS followed by 4% PFA. After post-fixation for 24 h, the brains were cryopreserved and cryosectioned at 30 µm with a Leica SM2010R microtome. For immunofluorescence, the brain slices were washed 3X in PBS, permeabilized in 0.5% Triton X-100-PBS solution for 0.5 hour, and then blocked in blocking buffer (10% normal donkey serum/NDS, 0.1% Triton X-100-PBS solution) for 1 hour. Following the blocking procedure, slices were incubated with primary antibodies diluted in the blocking buffer (Table 4) overnight at 4°C with agitation. The next day, slices were washed 3X in PBS and then incubated with specific fluorescent dye conjugated secondary antibodies diluted in PBS (Table 4) for 2 hours. Finally, they were mounted with Vectashield antifade mounting medium (Vector Laboratories: H-1000-10, Newark, CA, USA) on glass slides. Fluorescent images were acquired via an Olympus FV3000 confocal microscope. Images were then processed with ImageJ Fiji software. Pseudo-coloring was applied in certain images to improve visibility.

### 2.10 iDISCO

Following perfusion and post-fixation in 4% PFA, samples were rinsed with PBS 3X and stored in PBS with 0.02% NaN_3_ at 4LJC until the iDISCO protocol was performed. One hemisphere of each brain was used in iDISCO. Sample treatment, immunolabeling, and clearing were performed as described in the iDISCO protocol^14^ (available at idisco.info), which are briefly outlined below. Brains were kept in individual 5mL Eppendorf tubes (Hamburg, Germany) during the process, and gentle agitation was applied at each step unless otherwise noted.

#### Sample pretreatment

Brains were dehydrated serially in a methanol/ddH_2_O series: 20%, 40%, 60%, 80%, and 100% for 1 hour each. After the last methanol wash, samples were transferred to a fresh 100% methanol wash and kept at 4LJC for 1 hour. Brains were then incubated overnight in a 66% dichloromethane (DCM) / 33% methanol solution. The next day, brains were washed 2X in 100% methanol and then kept at 4LJC. Following this, brains were bleached in fresh, chilled 5% H_2_O_2_ in methanol overnight at 4LJC. The next day, brains were rehydrated with a methanol/ddH_2_O series: 80%, 60%, 40%, 20%, PBS for 1 hour each. Finally, brains were washed 2X for 1 hour each in 0.2% Triton X-100 in PBS (PTx.2).

#### Immunolabeling

Brains were permeabilized in PTx.2 / 2.3% (w/v) glycine / 20% DMSO for 48 hours at 37LJC and then blocked in PTx.2 / 6% donkey serum / 10% DMSO for 48 hours at 37LJC, followed by incubation with the primary antibody: rabbit anti-c-fos antibody (Synaptic Systems, Cat No: 226003; Göttingen, Germany) diluted (1:200) in PBS + 0.2% Tween-20 + 1% (w/v) heparin (PTwH) / 5% DMSO / 3% donkey serum for 120 hours at 37LJC. Brains were then washed in PTwH 5X for 1 hour each and incubated with the secondary antibody: Alexa Fluor 647 donkey anti-rabbit antibody (Jackson ImmunoResearch, Cat No: 711-605-152) diluted (1:200) in PTwH / 3% donkey serum for 120 hours at 37LJC. After the secondary incubation, brains were again washed in PTwH 5X for 1 hour each.

#### Clearing

After immunolabeling, brains were dehydrated in a methanol/ddH_2_O series: 20%, 40%, 60%, 80%, and 100% for 1 hour each. Then, brains were incubated for 3 hours in the 66% DCM / 33% methanol solution, and washed 2X for 15 minutes each with 100% DCM. Finally, brains were incubated in dibenzyl ether (DBE) without shaking until they were imaged.

### 2.11 Light sheet microscopy and ClearMap

Cleared iDISCO brains were imaged on a LaVision UltraMicroscope II light sheet microscope (LaVision BioTec, Bielefeld, Germany) using an Olympus MVPLAPO 2X objective with a custom LaVision BioTec dipping cap. The microscope chamber was filled with DBE, and brains were imaged at 1.6X in the sagittal orientation (lateral side up) with a step-size of 3 µm using the continuous light sheet scanning method for the 640 nm channel (20 acquisitions per plane) and the 480 nm channel for autofluorescence (without horizontal scanning). Quantification of c-fos positive cells was performed with the open-source ClearMap software (https://github.com/ChristophKirst/ClearMap) on an Ubuntu 14 computer. Images were aligned to the Allen Brain Atlas, background was corrected, and cells were detected with a detection threshold of 20 voxels for cell size. Cell counts were compared using a 2-sample student t-test, assuming unequal variances.

### 2.12 Fiber photometry

The social interaction assay was performed in a camera-monitored open field arena (L x W x H: 50 x 25 x 25 cm) located in a dimly lit (∼20 lux), sound-attenuating box. The day before behavioral testing, mice were gently handled and acclimated to the arena. In the 5-HT biosensor experiment, the Amuza wireless fiber photometry system was used, and recordings were conducted with 470 nm LED excitation, 90 µW light power, and 100 Hz sampling rate. First, the mouse was mounted with a headstage, and placed in the arena equipped with a small, empty corner cage. Prior to recording, the mouse was allowed to acclimate to the arena for 20 min.

The social interaction assay was split into two phases: a 10-min baseline phase during which the mouse was free to explore, followed by a 10-min social phase, which began with a novel C57BL/6J mouse of the same age and sex being carefully placed in the corner cage. Videos of the experiments were recorded using Ethovision XT 14 software and scored manually offline. During the baseline phase, facing, sniffing, or climbing on the empty cage were scored as exploration. After introduction of the stranger mouse, all the behaviors specified above were scored as social interaction. Fiber photometry data were acquired using TeleFipho software (Amuza) and analyzed offline with MATLAB R2021b software (MathWorks, Natick, MA, USA).

First, to correct for photobleaching, we created a linear regression model, using the MATLAB *fitlm* function to fit the raw data. Second, we normalized each fluorescent intensity value to the corresponding value predicted by the linear regression model. Finally, we computed peri-event fluorescent changes (ΔF/F) between the last 10 s pre-event (baseline) and the first 10 s post-event, and compared area under the curve (AUC) in the peri-event plot during the first 10 s of the first social interaction versus empty cage investigation.

In the dopamine biosensor experiment, we utilized the same procedure except that a wired fiber photometry system, Neurophotometrics FP3001, was employed. Therefore, instead of being mounted with a headstage, the mouse was tethered with a low-autofluorescence bundle branching fiber-optic patch cord connected to FP3001, and fiber photometry was recorded with interlaced 415 nm and 470 nm LED stimulation at 50 µW light power and 40 Hz sampling rate. All fiber photometry data were acquired using Bonsai software (Open Ephys, Atlanta, GA, USA) and analyzed offline with MATLAB. To normalize the raw data for photobleaching correction, we first fitted the isosbestic data (415 nm) to a two-term exponential model. Then, we applied the dopamine-dependent data (470 nm) to the isosbestic model. Finally, the dopamine-dependent fluorescent intensity values were normalized to the corresponding values predicted by the model. Peaks were identified using MATLAB *findpeaks* function. Frequency was defined as the number of peaks during a bout divided by the length of the bout. Latency was defined as the time from the start of the behavioral bout until the next fluorescence peak. Amplitudes were normalized to the minimum and maximum for each recording.

### 2.13 Two-photon imaging

AAVs expressing the GRAB_5-HT3.5_ biosensor (AAV9-hSyn-5-HT3.5) were stereotaxically injected into the DRN of C57BL/6J mice, as described above. Four weeks later, *ex vivo* two-photon fluorescent imaging was conducted, using an Olympus FVMPE RS multiphoton microscope paired with Mai Tai Si:Sapphire laser (Spectra-Physics, Mountain View, CA, USA). First, acute brain slices containing the DRN were prepared. Then, they were transferred to a recording chamber filled with aCSF saturated with 95% O_2_/5% CO_2_, flowing at the rate of 2 ml/min. The excitation wavelength was 920 nm, and fluorescence was detected with a 495-540-nm filter. Data were analyzed using ImageJ and MATLAB software.

### 2.14 Transmission electron microscopy

*Sert-cre* mice were stereotaxically injected with AAV8.2-ef1a-DIO-synaptophysin-eYFP in the DRN, and transmission electron microscopy was utilized to observe synaptic junctions between eYFP-labeled presynaptic part and dynorphin A-labeled post-synaptic part in the ventral NAcc. Briefly, we first performed immunohistochemical staining on free-floating 45 µm NAcc sections. They were incubated with chicken anti-GFP (1:100; Cat No: GFP-1010; Aves Labs, Davis, CA, USA) and rabbit anti-dynorphin A (1:100; Cat No: H-021-03; Phoenix Pharmaceuticals, Burlingame, CA, USA) primary antibodies for 72 h at 4°C, and then with 4 nm colloidal gold conjugated donkey anti-chicken (1:100; Cat No: 703-185-155; Jackson ImmunoResearch) and 12 nm colloidal gold conjugated donkey anti-rabbit (1:100; Cat No: 711-205-152; Jackson ImmunoResearch) secondary antibodies for 4 hours at room temperature. Following post-staining processing and dehydration^42^, the sections were embedded in resin. Then, serial ultrathin (50-100 nm) sections were made with a Leica EM UC7 ultramicrotome and mounted on mesh grids (4-6 sections/grid; 3.05 mm diameter). Finally, they were scanned under a 120 kV Hitachi HT7800 transmission electron microscope (Hitachi High-Tech, Tokyo, Japan).

### 2.15 Statistical analysis

Paired or unpaired student *t* tests were used for comparisons of two groups with one independent variable. One-way ANOVA was applied for comparisons of more than two groups with one independent variable and two-way or three-way ANOVAs were applied for data sets with more than one independent variable. If ANOVAs revealed significant main or interaction effects, *post hoc* Bonferroni tests were employed for pairwise comparisons. In the case of unequal variances, Welch’s correction was applied. Statistical significance was ascertained when p < 0.05. All the statistical analyses were performed using GraphPad Prism 9 software (Dotmatics, Boston, MA, USA). Data are expressed as mean ± SEM unless specified otherwise.

## Supporting information

Extended Dara Figures

## Declaration of Competing Interest

The authors declare that they have no known competing financial interests or personal relationships that could have influenced or appeared to influence this work.

### Acknowledgements

We thank Mackenzie McKnight, Gabrielle Bierlein-De La Rosa, Jeff Stolley, Yu Xu, and Suzanne Mason for their excellent technical assistance. Post-mortem brain tissue was provided by the Iowa Neurobank, and light sheet microscopy was made possible by the Iowa Neuroscience Institute Neural Circuits and Behavior Core with funding from the Roy J. and Lucille A. Carver Charitable Trust. We acknowledge the University of Iowa Central Microscopy Research Facility and its funding from the Office of the Vice President of Research for work that utilized the Transmission Electron Microscope (Hitachi HT7800). We also thank the University of Iowa Genome Editing Facility directed by William Paradee for generating Tph2-p2A-flpO mice for these studies, which is supported in part by grants from the NIH and the Roy J. and Lucille A. Carver College of Medicine. In addition, we wish to thank Norma Sinclair, Rongbin Guan, and Joanne Schwarting for their technical expertise in generating transgenic mice. Finally, we would like to thank Emmanuel Darq and Brigitte Kieffer (University of Strasbourg, Strasbourg, France) for providing AAV plasmids expressing scrambled and pdyn shRNA.

## Funding

This work was funded through NIH grants R00 AA024215 and R01 AA028931 as well as BBRF grant #27530 to C.A.M. K.M.K. was supported by T32 NS045549, T.J. was supported by T32 HL007638, and S.P. was supported by T32 GM067795.

## Respective Contributions

C.A.M., R.W., and N.B. wrote and edited the manuscript. R.W. and K.M.K. performed stereotaxic surgeries, ethanol administration in mice, behavioral experiments, and confocal imaging with assistance from S.P. and D.K. C.A.M. performed whole-cell patch clamp electrophysiological recordings and data analysis. K.M.K. conducted iDISCO, light sheet microscopy, and data analysis with ClearMap. K.M.K and N.B. performed FISH experiments and analysis. T.J. and R.W. performed fiber photometry experiments and analysis with assistance from D.K. S.G.P. performed TEM experiments and imaging. R.W. performed confocal microscopy to verify fiber optic placements as well as dlight1.1 and GRAB_5-HT3.5_ expression in the NAcc. K.M.K. performed two-photon imaging to verify the GRAB_5-HT3.5_ biosensor. Q.W. developed viral vectors for transsynaptic post-synaptic neuronal labeling. M.N. provided diagnostic information on human cases provided by the Iowa Neurobank and M.H. provided neuropathology expertise and assistance in sectioning human NAcc tissue for FISH analysis.

## Notes

### Competing Interest Statement

The authors have declared no competing interest.

