## Extended Dara Figures for "Alcohol inhibits sociability via serotonin inputs to the nucleus accumbens"

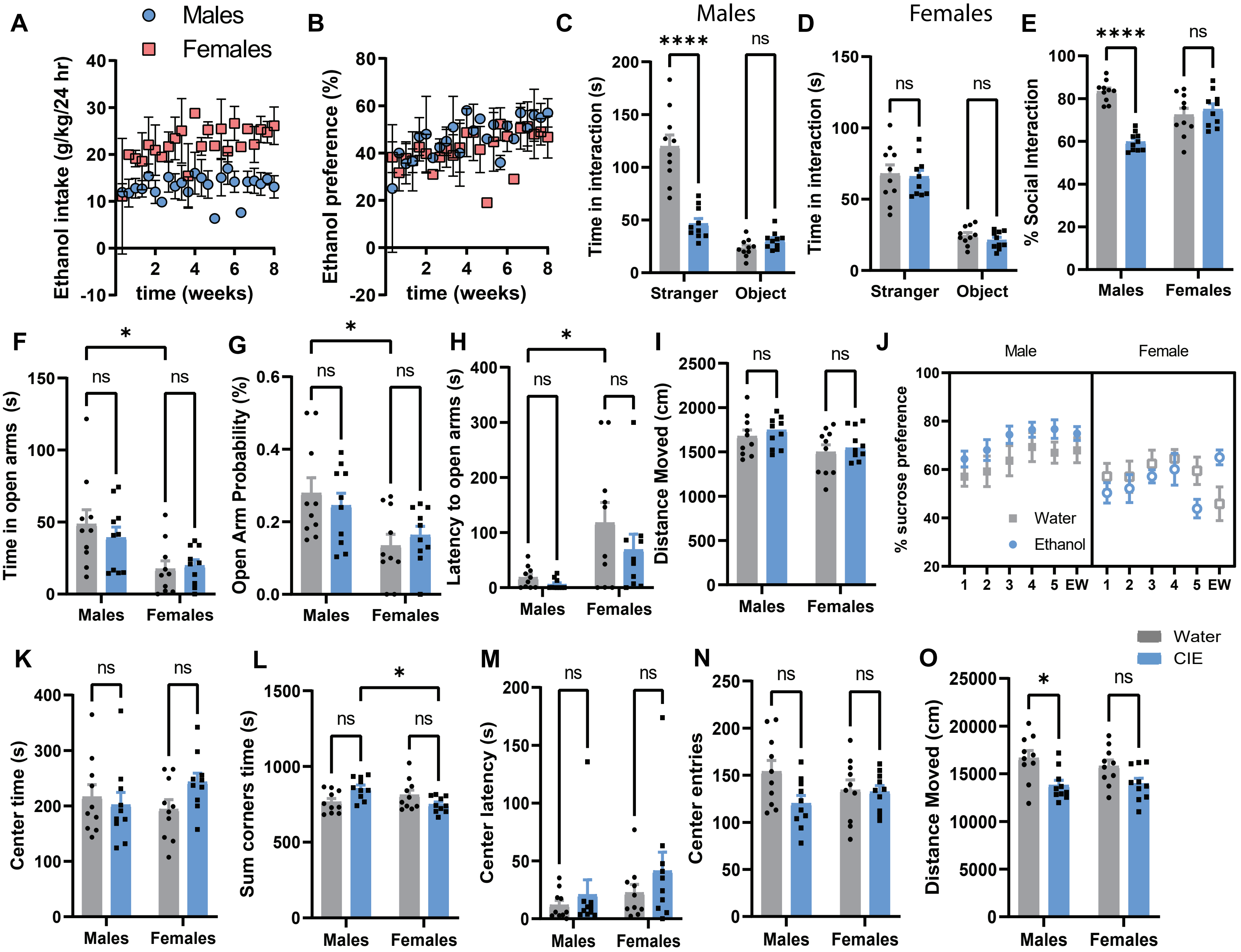


**Extended Data Figure 1: CIE promotes social deficits but not general anxiety- or depressive-like behaviors in male C57BL/6J mice.**

(A-B) Ethanol intake and preference over 8 weeks of CIE in male and female mice. (C-D) Time spent in social interaction or exploriation of an empty cage in male and female mice. (E) % time in social interaction relative to empty cage exploration in male and female mice. (F-I) Time in the open arms, open arm probability, latency to open arms and distance moved in the elevated plus maze (EPM). (J) % Sucrose preference. EW: ethanol withdrawal. (K-O) Center time, sum of corner time, center latency, center entries, and distance moved in the open field test. *p<0.05, ****p<0.0001, ns: non-significant.

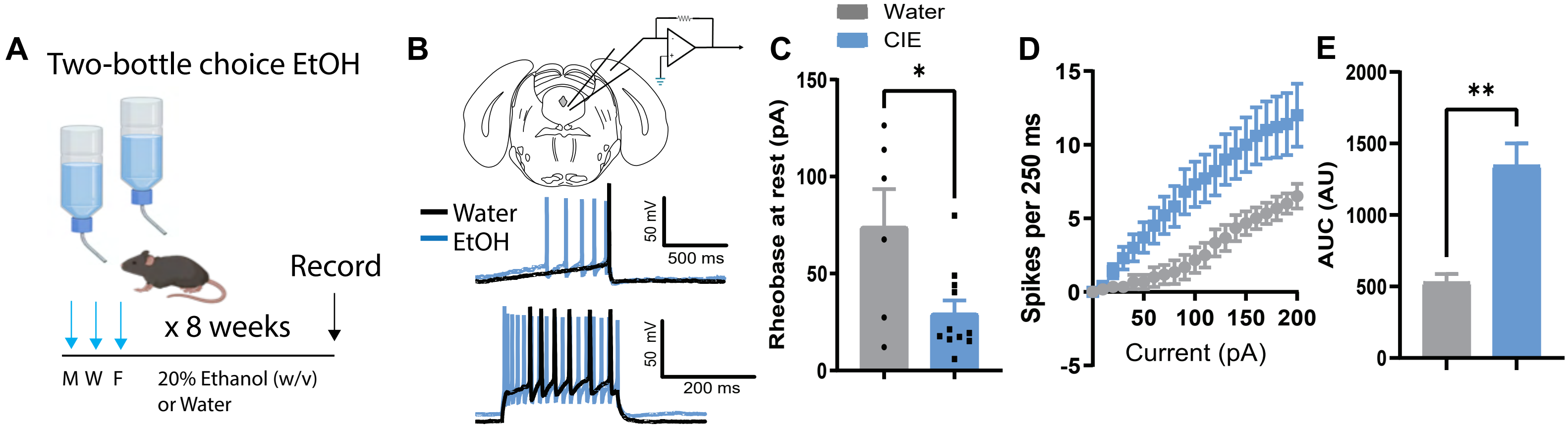


**Extended Data Figure 2: CIE enhances the intrinsic excitability of 5-HT$^{DRN}$ neurons.**

(A) Experimental schematic of CIE and whole-cell patch clamp recordings in DRN slices. (B) Representative traces depicting current ramp and current step (0-200 pA, 10 pA steps) protocols used to compute rheobase and action potential frequency. (C) Histogram of rheobase (action potential threshold), (D) current-induced spiking, and (E) area under the curve (AUC) of (D) in control and CIE mice. *p<0.05, **p<0.01, ns: non-significant. Parts of this figure were made using Biorender.

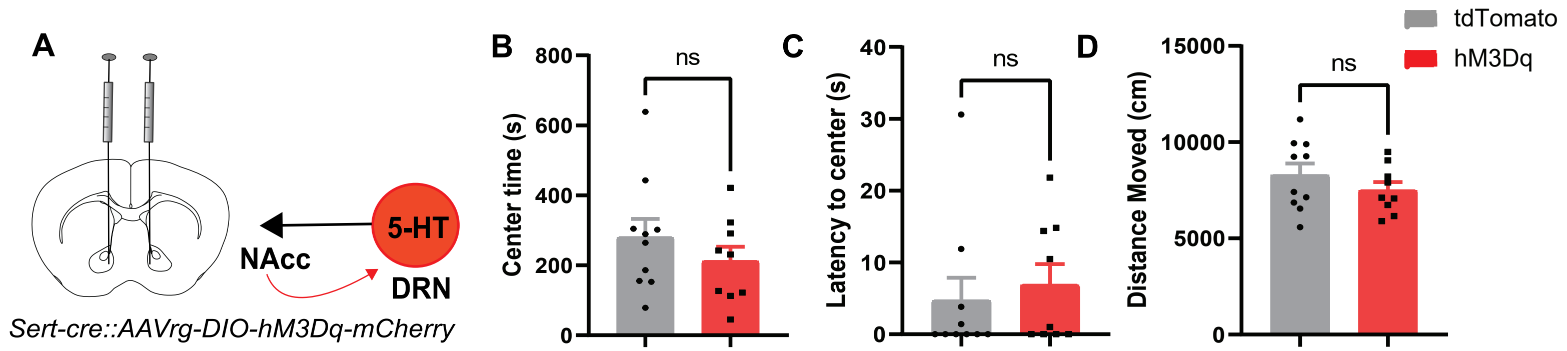


**Extended Data Figure 3: Chemogenetic activation of NAcc-projecting 5-HT neurons does not alter anxiety-like behavior or locomotor activity in the open field.**

(A) Schematic depicting stereotaxic injections of AAVrg-DIO-hM3Dq-mCherry into the NAcc of Sert-cre mice. (B-D) Center time, latency to center, and distance moved in the open field after chemogenetic stimulation of NAcc-projecting 5-HT neurons. ns: non-significant.

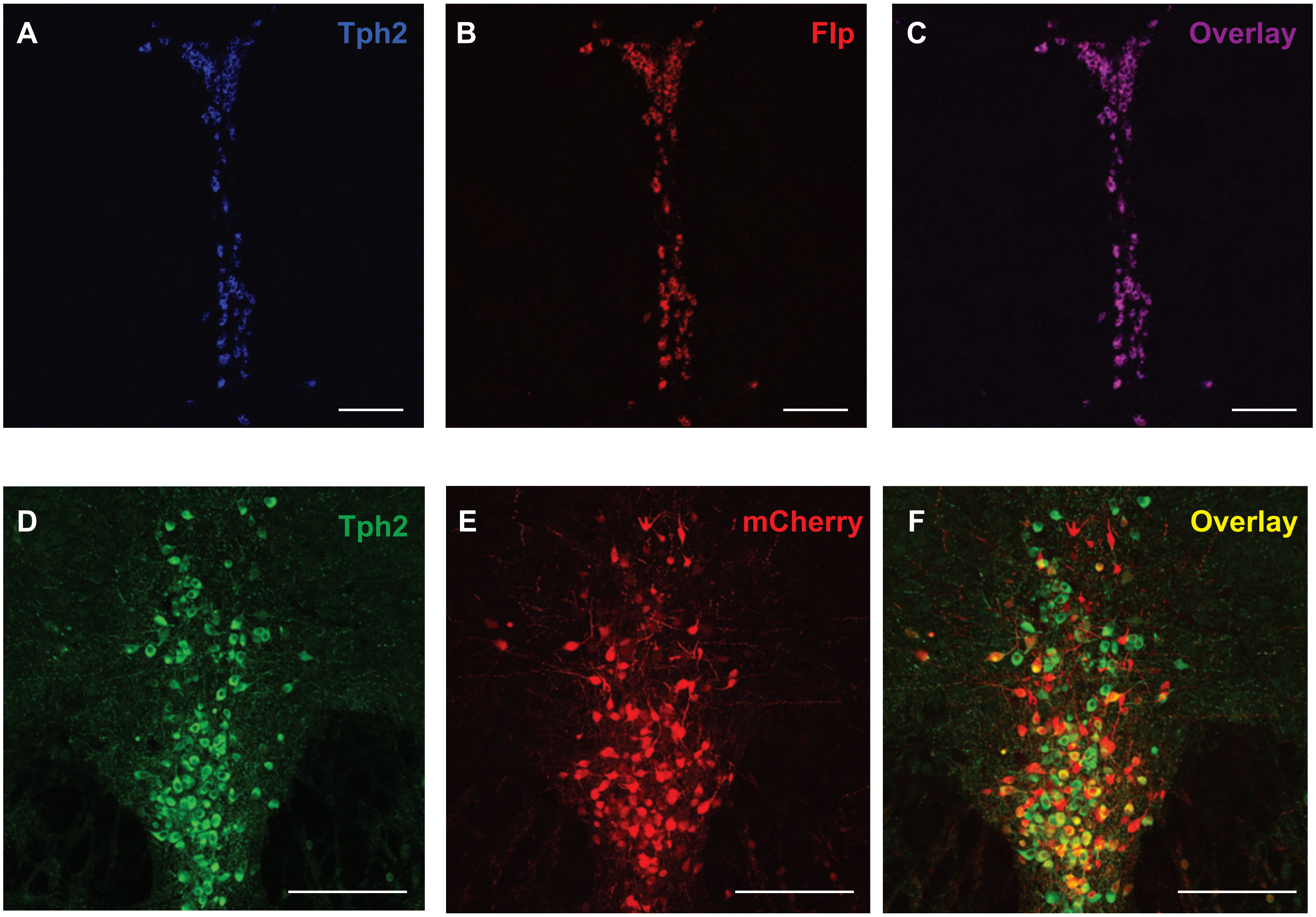


**Extended Data Figure 4: Generation and validation of *Tph2-p2A-flpO* mice.**

(A-C) Fluorescence in situ hybridization images depicting *Tph2*, *flp* and colocalization between the two in the DRN of *Tph2-p2A-flpO* mice. (D-F) Confocal images from *Tph2-p2A-flpO* mice stereotaxically injected with AAV5-ef1α-fDIO-mCherry in the DRN depicting Tph2, mCherry, and colocalization between the two in the DRN. Scale bars = 200 μm.

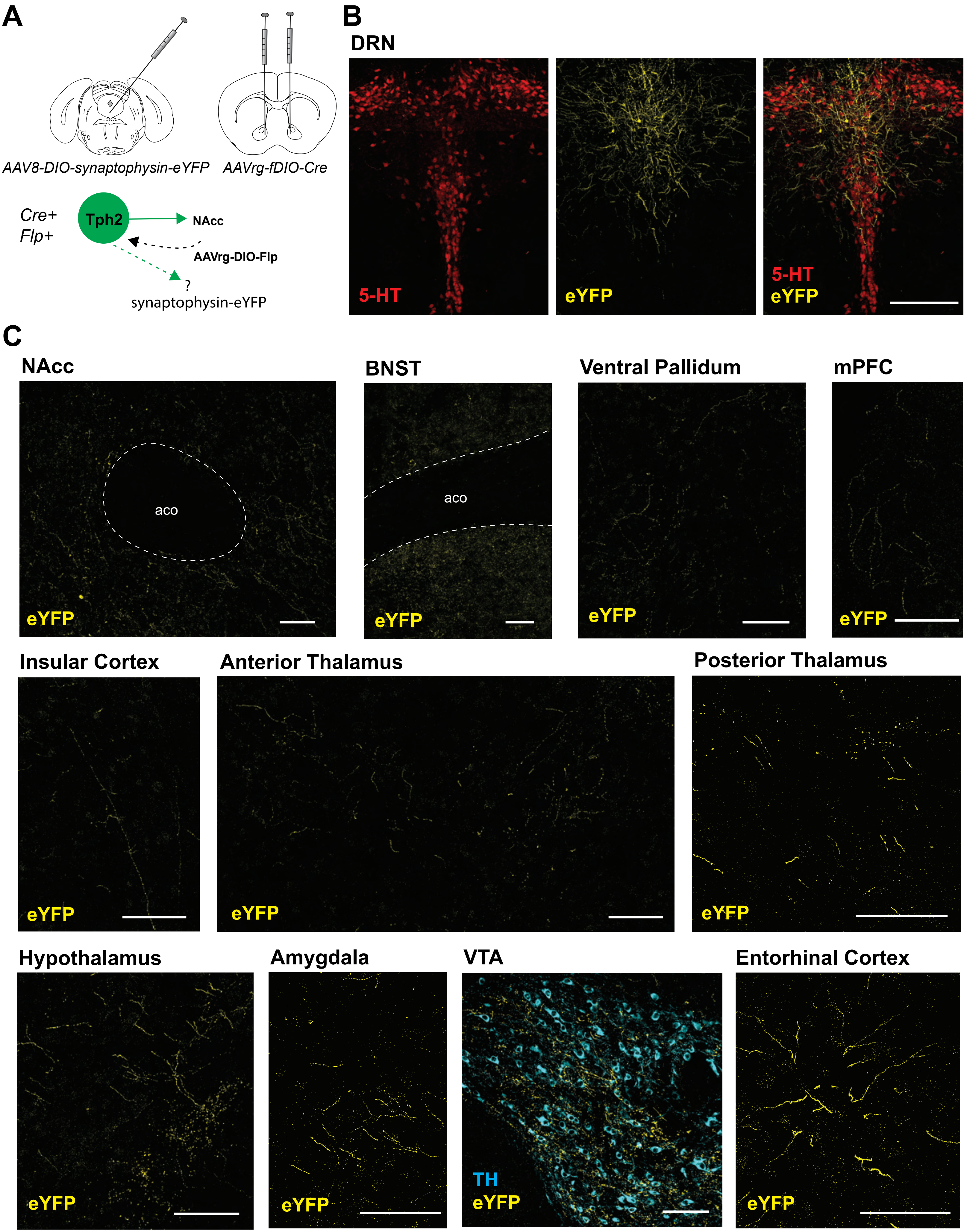
A
AAV8-DIO-synaptophysin-eYFP
AAVrg-fDIO-Cre
Cre+
Flp+
Tph2
NAcc
AAVrg-DIO-Flp
?
synaptophysin-eYFP
B
DRN
5-HT
eYFP
5-HT
eYFP
C
NAcc
aco
eYFP
BNST
aco
eYFP
Ventral Pallidum
eYFP
mPFC
eYFP
Insular Cortex
eYFP
Anterior Thalamus
eYFP
Posterior Thalamus
eYFP
Hypothalamus
eYFP
Amygdala
eYFP
VTA
TH
eYFP
Entorhinal Cortex
eYFP

**Extended Data Figure 5: 5-HT$^{DRN \rightarrow NAcc}$ neurons have collateral projections to other limbic and cortical structures.**

(A) Experimental schematic of viral targeting of Cre-dependent synaptophysin-eYFP to NAcc-projecting 5-HT$^{DRN}$ neurons in *Tph2-p2A-FlpO* mice. (B) Confocal images of synaptophysin-eYFP in the DRN. Scale bar = 300 µm (C) Confocal images of synaptophysin-eYFP in NAcc and other brain regions. Scale bars = 100 µm. Abbreviations: aco – anterior commissure, BNST – bed nucleus of the stria terminalis, mPFC – medial prefrontal cortex, VTA – ventral tegmental area, TH – tyrosine hydroxylase.

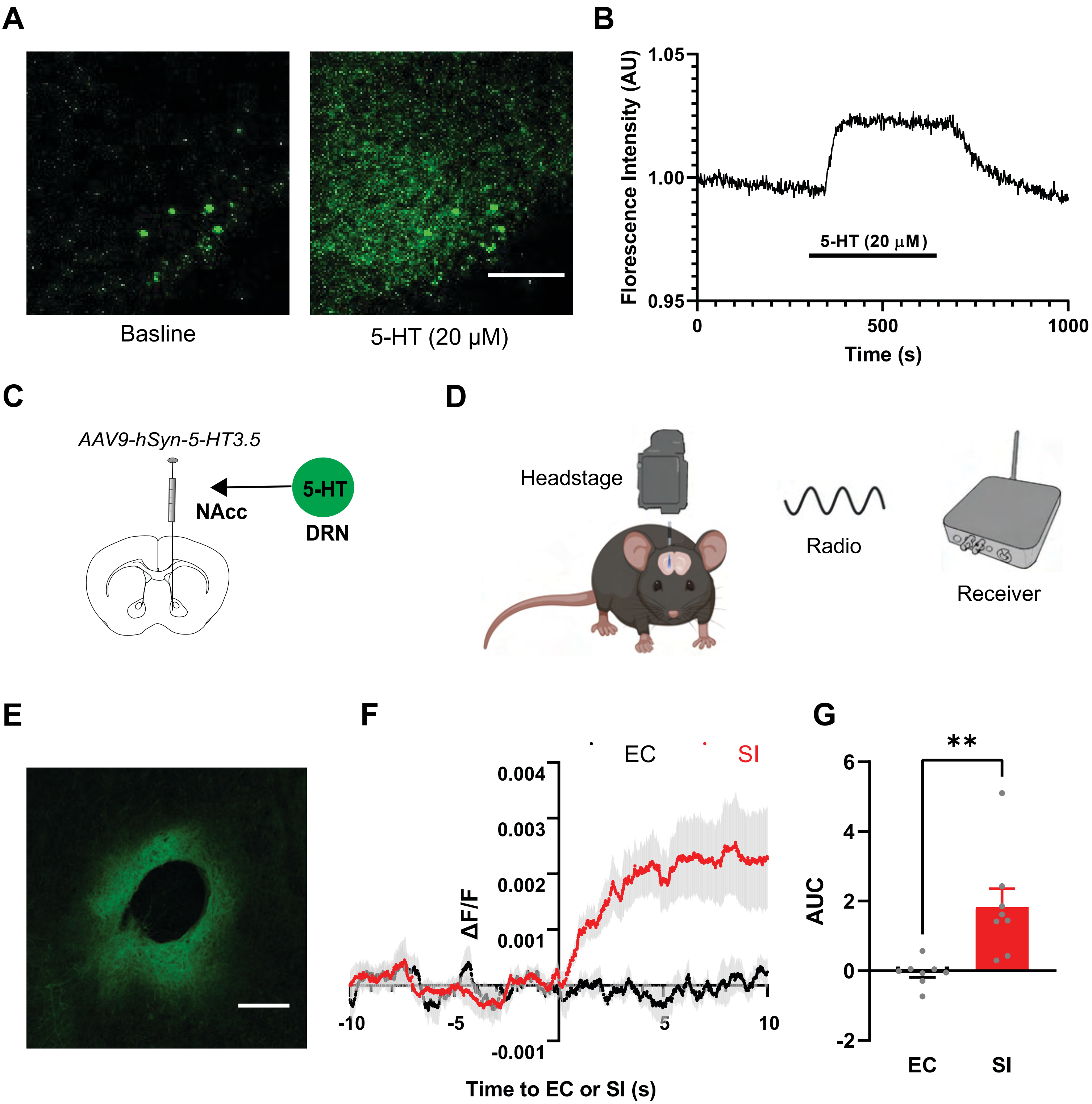


**Extended Data Figure 6: 5-HT release in the NAcc during social interaction.**

(A) Representative two-photon images showing an increase in fluorescent intensity, upon 5-HT (20 μM) application on an acute DRN slice expressing the $GRAB_{5\text{-}HT3.5}$ biosensor. Scale bar = 50 μm. (B) Experimental timeline of the 5-HT-induced increase in fluorescent intensity (AU: arbitrary unit), validating that changes in fluorescent intensity indicate changes in 5-HT levels. (C) Stereotaxic infusions of AAV expressing 5-HT biosensor $GRAB_{5\text{-}HT3.5}$ in the NAcc. (D) Experimental schematic of wireless fiber photometry recordings using Amuza TeleFipho system. (E) A representative confocal image of an NAcc slice expressing $GRAB_{5\text{-}HT3.5}$. Scale bar = 200 μm. (F) Peri-event plot showing changes in fluorescent intensity upon the first social interaction (SI) versus empty cage exploration (EC). Baseline: 10 s before the onset of the event (i.e., -10 ~ 0 s). Shade denotes standard error. (G) Histogram of area under the curve (AUC) computed from results between 0 and 10 s shown in (D). **p<0.01. *Parts of this figure were made using Biorender.*

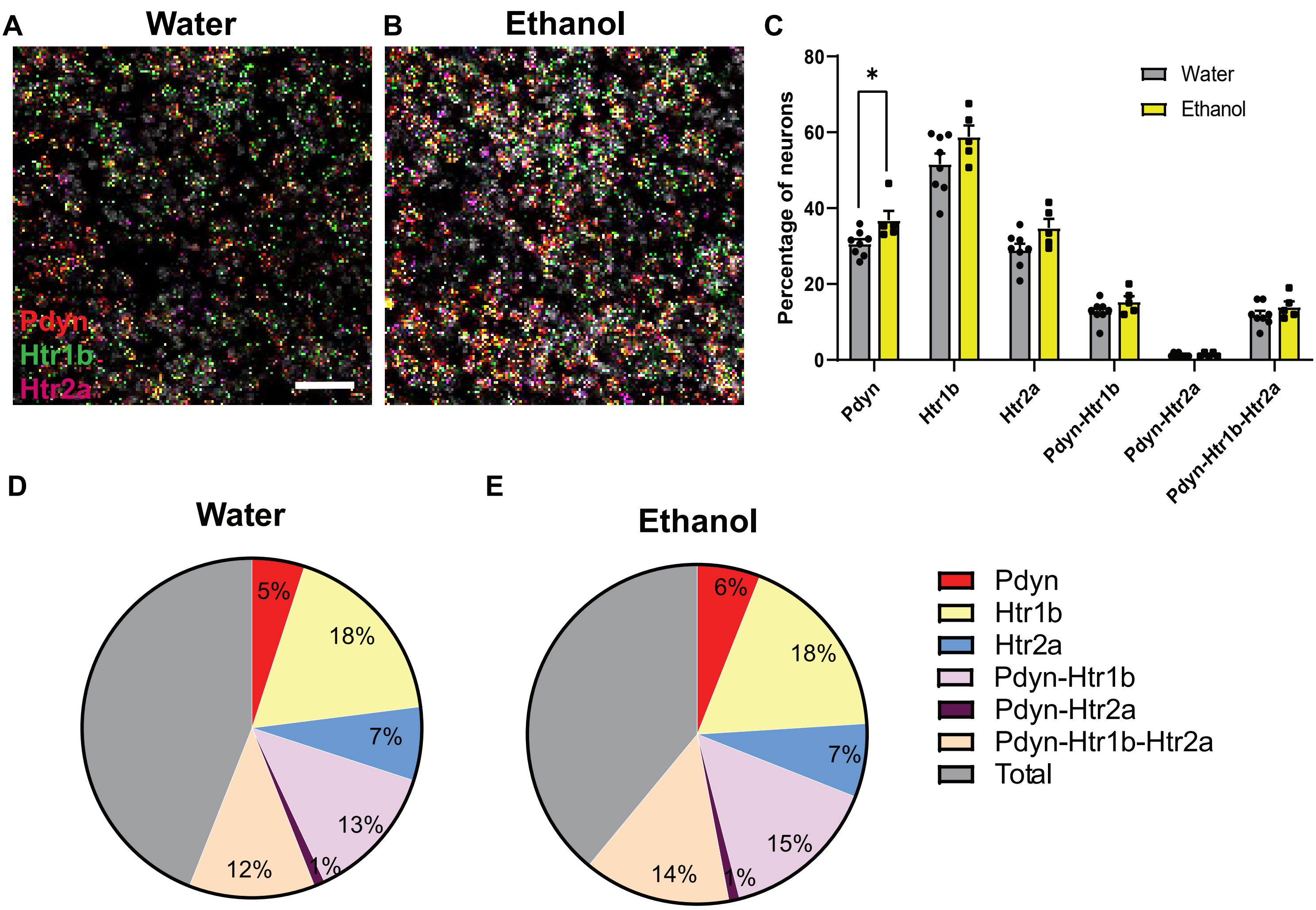


**Extended Data Figure 7: CIE does not alter 5-$HT_{2A}$ or 5-$HT_{1B}$ receptor expression in NAcc dynorphin neurons.**

(A-B) Fluorescence in situ hybridization images of *Pdyn*, *Htr1b,* and *Htr2a* receptors in the NAcc from control and CIE mice (scale bar = 50 µm). (C) Histogram of the percentages of total *Pdyn*, *Htr1b*, *Htr2a*; double positive for *Pdyn-Htr1b* and *Pdyn-Htr2a*; and triple positive for *Pdyn-Htr1b-Htr2a* neurons in the NAcc in control and CIE mice. (D-E) Pie chart depicting the percentages of neurons that express *Pdyn* only, *Htr1b* only, *Htr2a* only; double positive for *Pdyn-Htr1b* and *Pdyn Htr2a*; triple positive for *Pdyn-Htr1b-Htr2a*. *$p<0.05$
